# Munc18-1 catalyzes neuronal SNARE assembly by templating SNARE association

**DOI:** 10.1101/413724

**Authors:** Junyi Jiao, Mengze He, Sarah A. Port, Richard W. Baker, Yonggang Xu, Hong Qu, Yujian Xiong, Yukun Wang, Huaizhou Jin, Travis J. Eisemann, Frederick M. Hughson, Yongli Zhang

## Abstract

Sec1/Munc18-family (SM) proteins are required for SNARE-mediated membrane fusion, but their mechanism(s) of action remain controversial. Using single-molecule force spectroscopy, we found that the SM protein Munc18-1 catalyzes step-wise zippering of three synaptic SNAREs (syntaxin, VAMP2, and SNAP-25) into a four-helix bundle. Catalysis requires formation of an intermediate template complex in which Munc18-1 juxtaposes the N-terminal regions of the SNARE motifs of syntaxin and VAMP2, while keeping their C-terminal regions separated. Next, SNAP-25 binds the templated SNAREs to form a partially-zippered SNARE complex. Finally, full zippering displaces Munc18-1. Munc18-1 mutations modulate the stability of the template complex in a manner consistent with their effects on membrane fusion, indicating that chaperoned SNARE assembly is essential for exocytosis. Two other SM proteins, Munc18-3 and Vps33, similarly chaperone SNARE assembly via a template complex, suggesting that SM protein mechanism is conserved.

## Introduction

Cytosolic SM proteins and membrane-anchored SNARE proteins constitute the core machinery that mediates nearly all intracellular membrane fusion (Rizo and Sudhof, 2012; Sudhof and Rothman, 2009). In particular, the neuronal SM protein Munc18-1 and its cognate SNAREs syntaxin-1, SNAP-25, and VAMP2 (also called synaptobrevin) drive fusion of synaptic vesicles with the presynaptic plasma membrane (Sollner et al., 1993; Verhage et al., 2000). Fusion releases neurotransmitters into synaptic or neuromuscular junctions, controlling all thoughts and actions. Related SM proteins, Munc18-2 and Munc18-3, are required for cytotoxin release from lymphocytes to kill cancerous or infected cells (Cote et al., 2009) and for glucose uptake (Bryant and Gould, 2011), respectively. Consequently, dysfunctions of SM proteins are associated with neurological and immunological disorders, cancers, diabetes, and other diseases (Bryant and Gould, 2011; Cote et al., 2009; Stamberger et al., 2016).

SM proteins regulate the assembly of SNAREs into the membrane-bridging ‘trans-SNARE’ complexes required for membrane fusion (Figure 1) (Baker and Hughson, 2016; Brunger et al., 2018; Gao et al., 2012; Rizo and Sudhof, 2012; Shen et al., 2007; Sudhof and Rothman, 2009; Sutton et al., 1998). Most SNAREs contain a C-terminal transmembrane anchor, an adjacent SNARE motif, and an N-terminal regulatory domain (NRD). SNARE motifs are 60-70 residues in length, with either glutamine (Q-SNAREs) or arginine (R-SNAREs) residues at a key central position (Fasshauer et al., 1998). SNARE motifs in isolation are intrinsically disordered. By contrast, they are α-helical in fusion-competent SNARE complexes, with three Q-SNARE motifs (designated Qa, Qb, and Qc) and one R-SNARE motif combining to form a parallel four-helix bundle (Sutton et al., 1998). Despite its apparent simplicity, however, the physiological pathway(s) of SNARE assembly have remained enigmatic, as have the specific role(s) of SM proteins (Baker et al., 2015; Jakhanwal et al., 2017; Lai et al., 2017; Ma et al., 2013; Ma et al., 2015; Rizo and Sudhof, 2012; Shen et al., 2007; Wickner, 2010; Zhang et al., 2015; Zhou et al., 2013).

**Figure 1.**
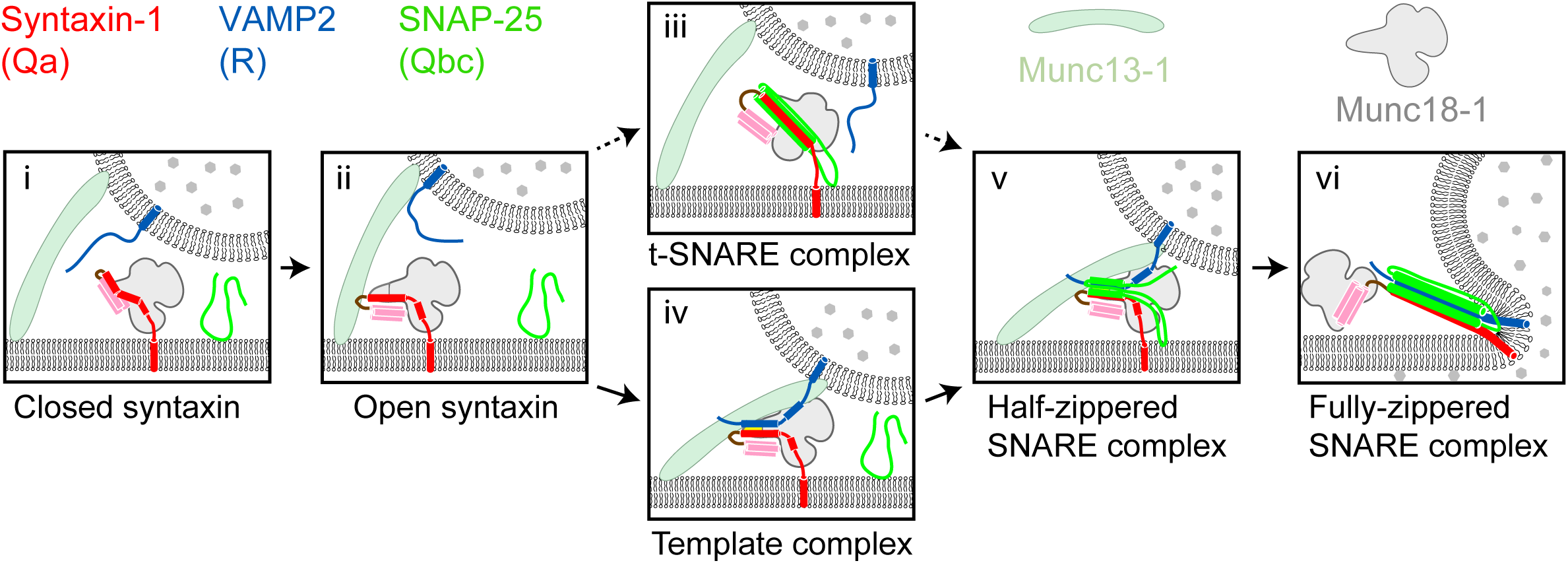
Two potential pathways for Munc18-1-regulated neuronal SNARE assembly. (i) Munc18-1 first serves as a syntaxin chaperone and binds syntaxin to inhibit its association with other SNAREs. (ii) Closed syntaxin is opened by Munc13-1, a large multifunctional protein that also helps tether vesicles to the plasma membrane and binds, albeit with low affinity, both syntaxin and VAMP2. (iii) Open syntaxin may bind SNAP-25 to form a syntaxin:SNAP-25 or a Munc18-1:syntaxin:SNAP-25 complex. (iv) Alternatively, open syntaxin may bind VAMP2 to form a Munc18-1:syntaxin:VAMP2 template complex, as proposed here. Both complexes, (iii) and (iv), have been proposed to be ‘activated’ for SNARE assembly. (v) and (vi) Other factors such as synaptotagmin (not shown) target the half-zippered SNARE complex to enable calcium-triggered further SNARE zippering and vesicle fusion.

SNARE assembly has long been thought to begin with the formation of a t-SNARE complex among the SNAREs – usually Qa, Qb, and Qc – residing on the target membrane (Weber et al., 1998) (Figure 1). According to this view, the neuronal SNAREs syntaxin (Qa-SNARE) and SNAP-25 (Qbc-SNARE, a single protein containing both Qb and Qc SNARE motifs) assemble on the presynaptic plasma membrane, forming a t-SNARE complex that subsequently binds to the synaptic vesicle R-SNARE VAMP2 (Jakhanwal et al., 2017; Pobbati et al., 2006; Shen et al., 2007; Weber et al., 1998; Zhang et al., 2016a). Recent reports have, however, raised doubts about this order of events. In vitro reconstitution experiments suggested that neuronal SNARE assembly begins with a complex between Munc18-1 and syntaxin, requires Munc13-1, and may not involve a syntaxin:SNAP-25 complex (Ma et al., 2013) (Figure 1). Crystal structures of the SM protein Vps33 bound to its cognate Qa- and R-SNARE implied that the SM protein functions as a template, orienting and aligning the two SNARE motifs for further assembly (Baker et al., 2015). Thus the Qa- and R-SNAREs might be the first to assemble, and only on the surface of an SM template.

Previously, we developed a single-molecule approach based on optical tweezers to dissect SNARE assembly at high spatiotemporal resolution (Gao et al., 2012; Ma et al., 2015; Zhang et al., 2016a; Zorman et al., 2014). Using this method, we measured the folding energy and kinetics of various SNARE complexes. Here, we extend the method to observe SM-mediated SNARE assembly. We detected three template complexes, each of them comprising an SM protein (Munc18-1, Munc18-3, or Vps33) bound to its cognate Qa- and R-SNAREs, and characterized the neuronal template complex in detail using a large panel of mutant proteins. Our results imply that the neuronal template complex is an on-pathway, rate-limiting intermediate in vitro and in vivo. They further suggest that phosphorylation of Munc18-1 can modulate the efficiency of neurotransmitter release by affecting the stability of the template complex. More broadly, our findings imply that membrane fusion in vivo may be controlled by SM proteins through their tunable catalytic activity as SNARE assembly chaperones.

## Results

### Munc18-1, syntaxin, and VAMP2 form a template complex

Previously, we found that the SM protein Vps33 forms binary complexes with the SNARE motifs of Vam3 (Qa-SNARE) and Nyv1 (R-SNARE), as well as a ternary ‘template complex’ containing all three proteins (Baker et al., 2015). Crystal structures of the two binary complexes revealed that the Qa-SNARE and the R-SNARE bind to adjacent sites on the SM protein and led to a model of the template complex in which the two SNARE motifs are ‘half-zippered’. An analogous template complex might form during the assembly of the neurotransmitter release machinery (Sitarska et al., 2017), but direct evidence is lacking. To investigate further, we used an optical tweezers-based strategy to directly observe neuronal SNARE assembly and disassembly in the presence of Munc18-1. To mimic a trans-SNARE complex, pre-assembled SNAREs were attached via the C termini of the Qa- and R-SNARE motifs to beads (Gao et al., 2012). The same SNARE motifs were covalently linked near their N termini through an engineered disulfide bond to form a Qa-R-SNARE conjugate (Figure 2A & Figure 2-figure supplement 1). This tactic permitted us to conduct repeated rounds of force-induced unfolding/disassembly (‘pulling’) and potential refolding/assembly (‘relaxation’) in a single experiment.

**Figure 2.**
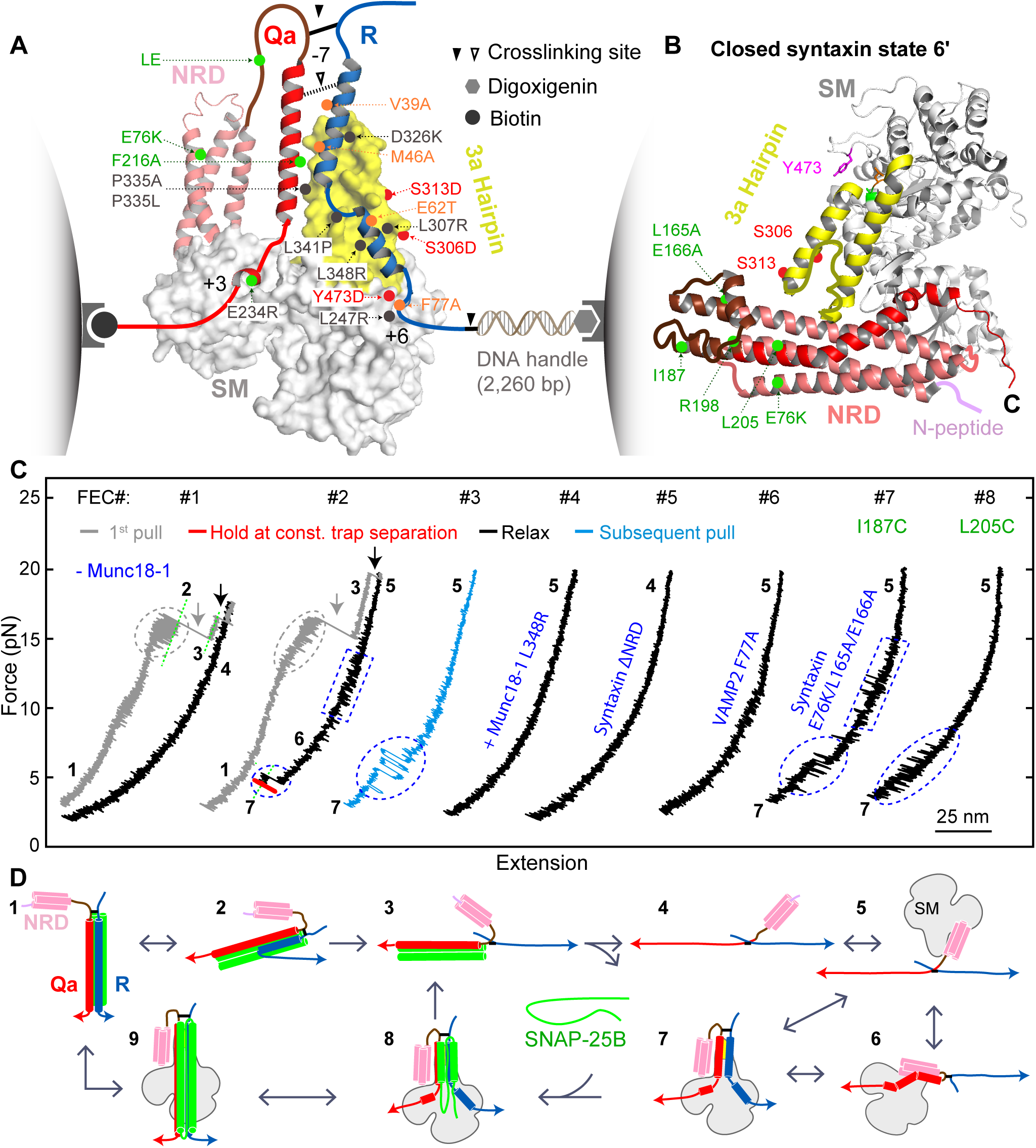
Single-molecule manipulation based on optical tweezers revealed a ternary template complex. (A) Experimental setup and structural model of the template complex. Some key mutations tested in this study are indicated by dots: red (phosphomimetic mutations) or gray (others) for Munc18-1, green for syntaxin, and orange for VAMP2. The helical hairpin of Munc18-1 domain 3a is highlighted in yellow. The NRD of syntaxin comprises an N-peptide (a.a. 1-26, see B), a three-helical Habc domain (27-146, deep salmon), and a linker region (147-199, brown). The structural model of the template complex is derived from a similar model of Vps33:Vam3:Nyv1 (Baker et al., 2015) by extending the N-terminal helix of the R-SNARE to −7 layer, as justified herein. The NRD stabilizes the template complex, but its positioning in this model is arbitrary. See also Figure 2-figure supplement 1. (B) Crystal structure of closed syntaxin bound to Munc18-1 (PDB ID 3C98) (Misura et al., 2000). Highlighted are crosslinking sites (I187, R198, and L205), sites of mutations used to destabilize closed syntaxin (E76K, L165A, and E166A, green dots), and sites of phosphomimetic mutations (red dots). (C) Force-extension curves (FECs) obtained in the absence (#1) or presence (other FECs) of Munc18-1 in solution. Throughout the figures, all FECs are color coded in the same fashion: gray for pulling the initial purified SNARE complex, blue for subsequent pulls, black for relaxation, and red for holding the Qa-R SNAREs at constant force. The states associated with different extensions (marked by green dashed lines as needed) are numbered as in Figure 2D. CTD transitions are indicated by gray ovals, NTD unfolding by gray arrows, t-SNARE unfolding by black arrows, syntaxin transitions by blue rectangles, and template complex transitions by blue ovals. See also Figure 2-figure supplement 2. (D) Schematic diagrams of different states: 1, fully assembled SNARE complex; 2, half-zippered SNARE bundle; 3, unzipped t-SNARE complex; 4, fully unfolded SNARE motifs; 5, unfolded SNARE motifs with Munc18-1 bound; 6, partially closed syntaxin; 7, template complex; 8, Munc18-1-stabilized partially-zippered SNARE complex containing SNAP-25; and 9, Munc18-1-bound assembled SNARE complex. The states are numbered according to the same convention throughout the text and figures.

Munc18-1 binds both the Qa-SNARE syntaxin (with nanomolar affinity) and the R-SNARE VAMP2 (with micromolar affinity) (Burkhardt et al., 2008; Misura et al., 2000; Parisotto et al., 2014; Sitarska et al., 2017). Formation of a ternary template complex has not, however, been reported. This is presumably because Munc18-1 and syntaxin, in their high-affinity complex, both adopt conformations that preclude VAMP2 binding (Baker et al., 2015; Misura et al., 2000; Sitarska et al., 2017) (Figure 2B). In particular, the SNARE motif and NRD of syntaxin interact to create an autoinhibited or ‘closed’ conformation (Misura et al., 2000). Opening syntaxin, and thereby permitting SNARE assembly, requires Munc13-1 for a mechanism that remains controversial (Ma et al., 2011; Ma et al., 2013; Wang et al., 2017; Yang et al., 2015). To bypass the requirement for Munc13-1 in our single-molecule experiments, we attempted to destabilize the closed conformation of syntaxin without abolishing its interactions with Munc18-1 or the other SNAREs. Among the strategies we evaluated, the simplest was to form the Qa-R-SNARE conjugate by crosslinking syntaxin R198C and VAMP2 N29C (Figure 2A, solid arrowhead; Figure 2-figure supplement 1). In closed syntaxin, residue 198 is buried against the NRD (Figure 2B). As shown below, involving this residue in a disulfide bond destabilized and partially opened Munc18-bound syntaxin, presumably via localized unfolding.

We began by pulling the fully folded neuronal SNARE complex, containing crosslinked syntaxin and VAMP2 as well as SNAP-25B, in the absence of Munc18-1. The resulting force-extension curve (FEC) revealed that, as expected based on our previous work (Gao et al., 2012; Ma et al., 2015), the SNARE complex disassembled in at least three steps (Figure 2C, FEC #1, gray curve). These force-induced disassembly steps are schematically depicted in Video 1 and in Figure 2D as transitions from states 1↔2→3→4. 1↔2 represents reversible unfolding of the C-terminal half of the VAMP2 SNARE motif (CTD; Figure 2C, gray oval), 2→3 represents irreversible unfolding of the N-terminal half of the VAMP2 SNARE motif (NTD; gray arrow), and 3→4 represents irreversible unfolding of the syntaxin SNARE motif (black arrow). 3→4 was accompanied by release of SNAP-25B. Relaxing the resulting Qa-R-SNARE conjugate revealed a featureless FEC, as expected for an unfolded polypeptide (Figure 2C, FEC #1, black curve) (Gao et al., 2012; Ma et al., 2015).

We next asked whether our single-molecule assay could be used to detect and characterize the predicted template complex (Figure 2A). The addition of 2 μM Munc18-1 had little effect on the unfolding pathway of the initial syntaxin/VAMP2/SNAP-25B complex (Figure 2C, compare gray curves in FEC #1 and #2; Video 1). However, the presence of Munc18-1 had a striking effect on the FEC of the remaining Qa-R-SNARE conjugate. Specifically, relaxing (Figure 2C, #2, black trace) and then pulling (Figure 2C, #3, blue trace) the Qa-R-SNARE conjugate revealed two Munc18-1-dependent features (Figure 2-figure supplement 2). In about 40% of the FECs, we observed a small flickering signal at 10-15 pN (Figure 2C, #2 in blue rectangle; Figure 2-figure supplement 3). We attribute this transition (5↔6) to the reversible folding/unfolding of the partially closed syntaxin conformation induced by Munc18-1 (state 6 in Figure 2D). More importantly, in about 50% of the FECs, we observed prominent flickering signals at 3-7 pN (Figure 2C, #2-3, blue ovals). As described in detail below, extensive evidence supports the conclusion that this transition (6↔7) results from the reversible, cooperative formation and unfolding of the predicted template complex (Figure 2A; state 7 in Figure 2D). For example, the probability of observing the 6↔7 transition was greatly reduced when either the Munc18-1:VAMP2 interaction or the Munc18-1:syntaxin interaction was abrogated (Burkhardt et al., 2008; Parisotto et al., 2014) (Figure 2C, #4-5; Table 1).

**Table 1.**
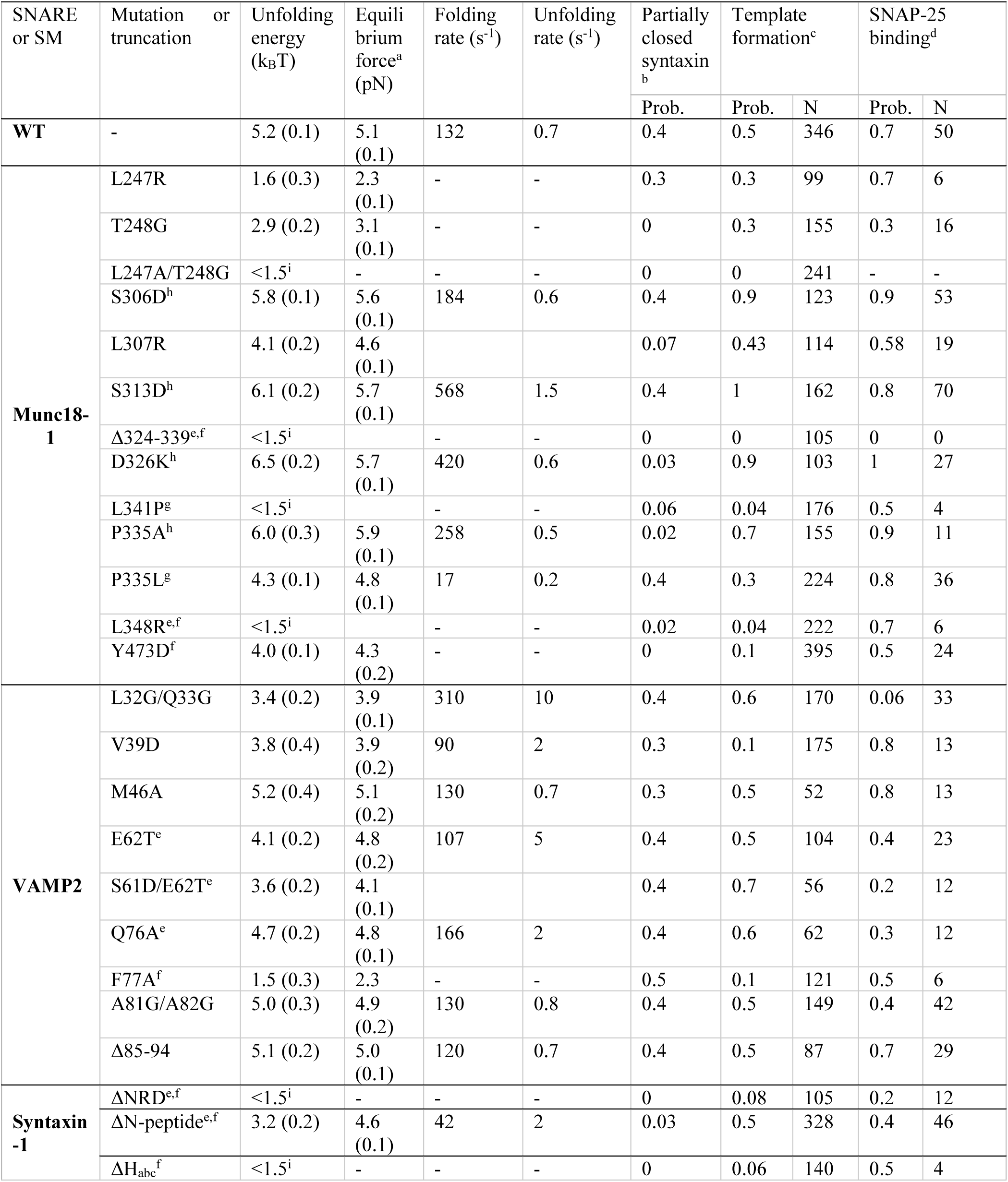
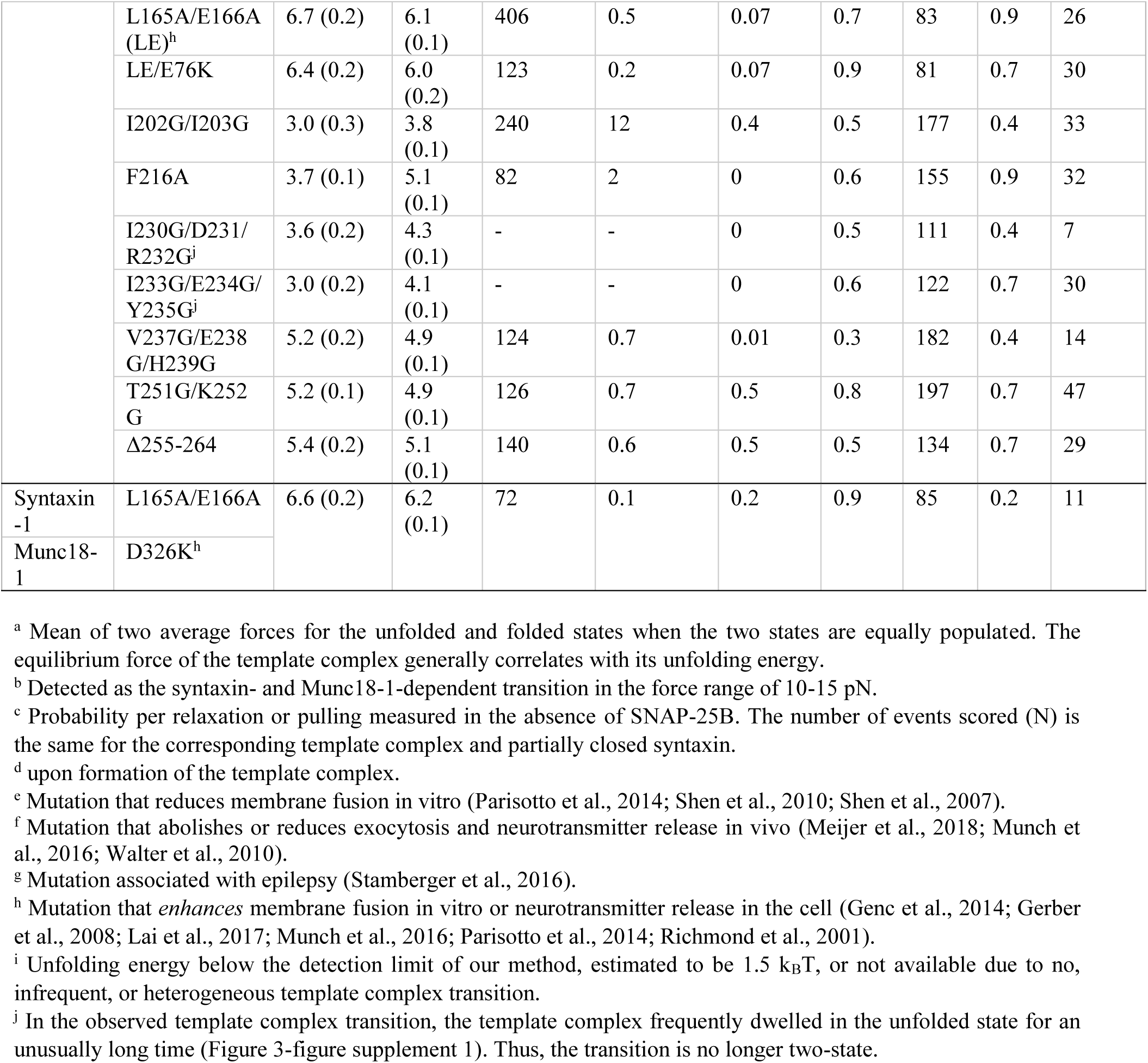
Properties of the neuronal template complex. The number in parenthesis is the standard error of the mean.

### Stability and conformation of the template complex

To examine the stability and folding/unfolding kinetics of the template complex, we monitored the 6↔7 transition over a range of constant mean forces (Figure 3A-C; Video 2). Detailed analyses of the extension trajectories (Figure 3A) revealed the force-dependent unfolding probability and transition rate of the template complex (Figure 3C) and, by extrapolation to zero force (Gao et al., 2012; Rebane et al., 2016), its unfolding energy (5.2 ± 0.1 kBT or 3.1 ± 0.1 kcal/mol; mean ± SEM) and lifetime (1.4 s) (Figure 3B). Comparable analysis of the folding/unfolding of the partially closed syntaxin (5↔6) allowed us to estimate its unfolding energy as well (2.6 ± 0.2 kBT; Figure 2-figure supplement 4). The total extension change associated with these transitions (5→6→7) is consistent with a structural model of the template complex based on the crystal structures of Vps33:Nyv1 and Vps33:Vam3 (Baker et al., 2015) (Figure 2A). Importantly, the same template complex was observed when we used an alternative Qa-R-SNARE crosslinking site at syntaxin I187C and VAMP2 N29C, but only in conjunction with additional NRD mutations E76K, L165A, and E166A to destabilize the closed conformation of syntaxin (Figure 2C, #7; Figure 2B; Figure 2-figure supplement 5-7). Thus, observation of the template complex was independent of the crosslinking site, requiring only that the closed conformation be destabilized (Hu et al., 2011).

**Figure 3.**
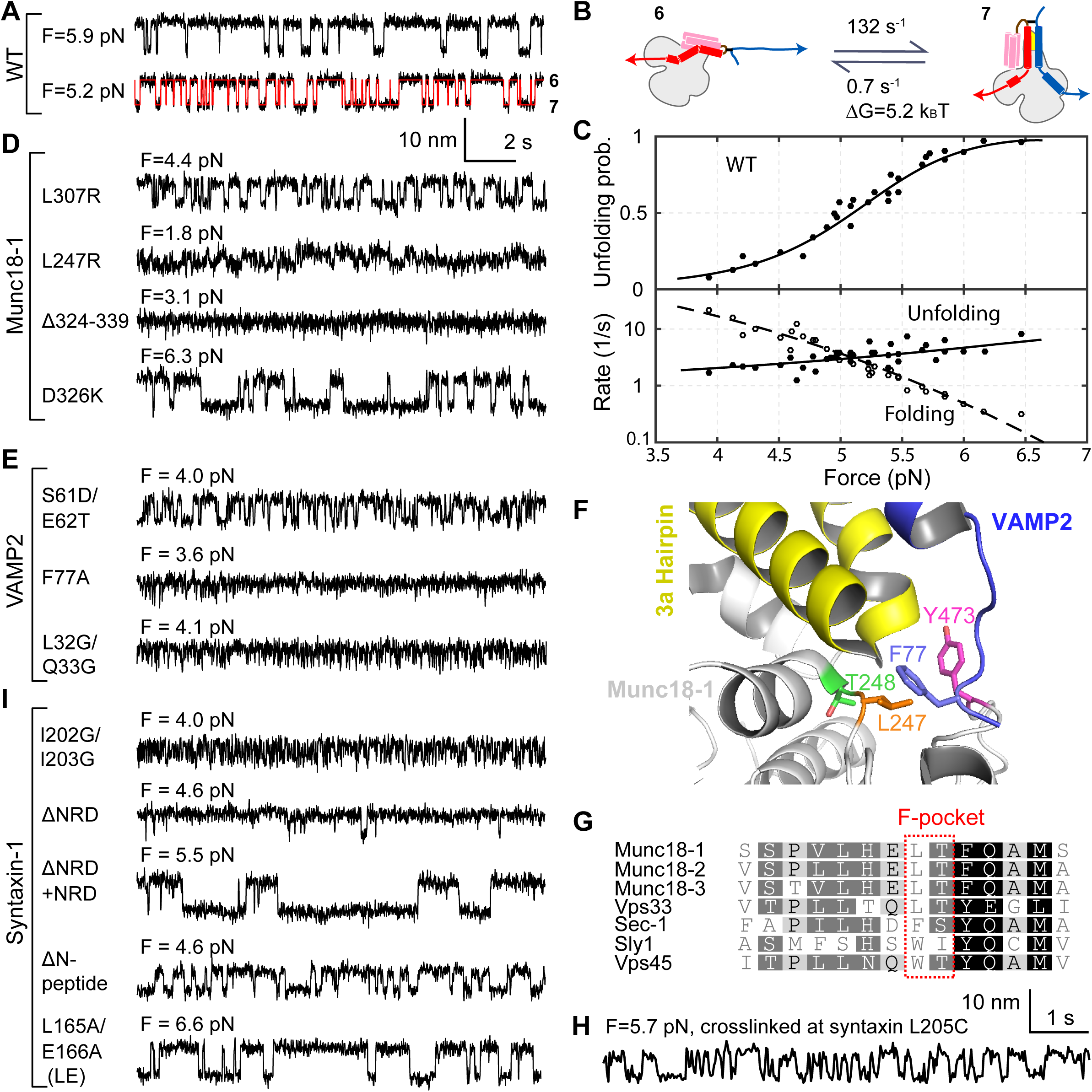
Stability, conformation, and folding kinetics of the template complex. (A, D, E, I) Extension-time trajectories at constant mean forces with the WT template complex or its variants containing indicated mutations in Munc18-1 (D), VAMP2 (E), or syntaxin (I). The red trace in A shows an exemplary idealized trajectory derived from hidden Markov modeling. Trajectories in A, D, E, and I share the same scale bars. See also Figure 3-figure supplement 1. (B)Diagram illustrating the transition between the partially closed syntaxin state (state 6 in Figure 2D) and the template complex state (state 7); rates and energies are derived from panel C. (C) Force-dependent unfolding probabilities (top) and transition rates (bottom). Best model fits (solid and dashed curves) reveal the stability and folding and unfolding rates of the template complex at zero force (Figure 4 and Table 1). (F) Structural model of VAMP2 F77 anchored in the F-pocket in Munc18-1 composed of L247 and T248, which is covered by Y473. The model is derived by superimposing the structures of Munc18-1:syntaxin (Figure 2B; 3C98) and Vps33:Nyv1 (5BV0). (G) Sequence alignment showing F-pocket sequence conservation among SM proteins. (H) Extension-time trajectory of the WT template complex at 5.7 pN. The Qa-R SNAREs were crosslinked between syntaxin L205C and VAMP2 Q36C (Figure 2A, open arrowhead). See also Figure 2-figure supplement 1.

We used a battery of mutant proteins to test our structural model of the template complex in greater detail (Figure 2A). A salient feature of the model is the pivotal role played by a pair of α-helices (a.a. 298-359) within domain 3a of Munc18-1 (Baker et al., 2015; Sitarska et al., 2017) (yellow in Figure 2A,B). These α-helices form an extended helical hairpin that interacts extensively with the NTD of syntaxin and with both the NTD and the CTD of VAMP2 (Figure 2A & Figure 2-figure supplement 1). Many domain 3a mutations within (L307R, P335L, L341P, L348R) or adjacent (L247R, T248G) to the helical hairpin destabilized the template complex (Figures 3D, 4, & Figure 3-figure supplement 1; Table 1 and references therein). An internal deletion that removes the distal portion of the helical hairpin (Munc18-1 Δ324-339) abolished formation of the template complex altogether. Notably, two helical hairpin mutations – D326K and P335A – actually stabilized the template complex; both of these mutations are associated with enhanced Munc18-1 function in vitro and in vivo (Munch et al., 2016; Parisotto et al., 2014; Sitarska et al., 2017). Three phosphomimetic mutations (S306D, S313D, and Y347D) are discussed later. None of the mutations we tested had a significant effect on the overall structure of Munc18-1 as judged by circular dichroism (Figure 3-figure supplement 2). Overall, the consequences of Munc18-1 mutations are consistent with our structural model.

Reciprocally, we investigated the impact of SNARE motif mutations that appeared likely to affect the SNARE:Munc18-1 interface. Although VAMP2 M46A did not have a significant effect, the rest (syntaxin F216A, I230G/D231/R232G, and I233G/E234G/Y235G; VAMP2 S61D/E62T, E62T, Q76A, and F77A) all destabilized the template complex (Figures 3E & 4; Table 1; Figure 3-figure supplement 1). Interestingly, all three syntaxin mutations abolished the partially closed syntaxin (Table 1), implying that the template complex and the closed syntaxin share some interactions between syntaxin and Munc18-1. The VAMP2 residue Phe 77, located at the so-called +6 layer (Figure 2-figure supplement 1), appears to play an especially important role. In our model of the template complex, the side chain of Phe 77 inserts into a deep, hydrophobic pocket in domain 3a, with Leu 247 and Thr 248 residues at the bottom (Figure 3F). Phe 77 is highly conserved among R-SNAREs, whereas Leu 247 and Thr 248 are highly conserved among SM proteins (Figure 3G). Substituting Phe 77 with Ala dramatically reduced the formation probability of the template complex to 0.06 and its unfolding energy to 1.5 ± 0.3 kBT, the lower limit of our assay (Figure 2C, #6; Figure 3E). Similarly, Munc18-1 mutations in the hydrophobic pocket strongly impaired (for L247R or T248G) or totally abolished (for L247A and T248G together) formation of the template complex (Figure 4; Table 1). Taken together, our mutagenesis results confirm that the stability of the template complex depends on extensive interactions between Munc18-1 and the two SNARE motifs, including a key anchoring role for the +6 layer Phe of VAMP2.

**Figure 4.**
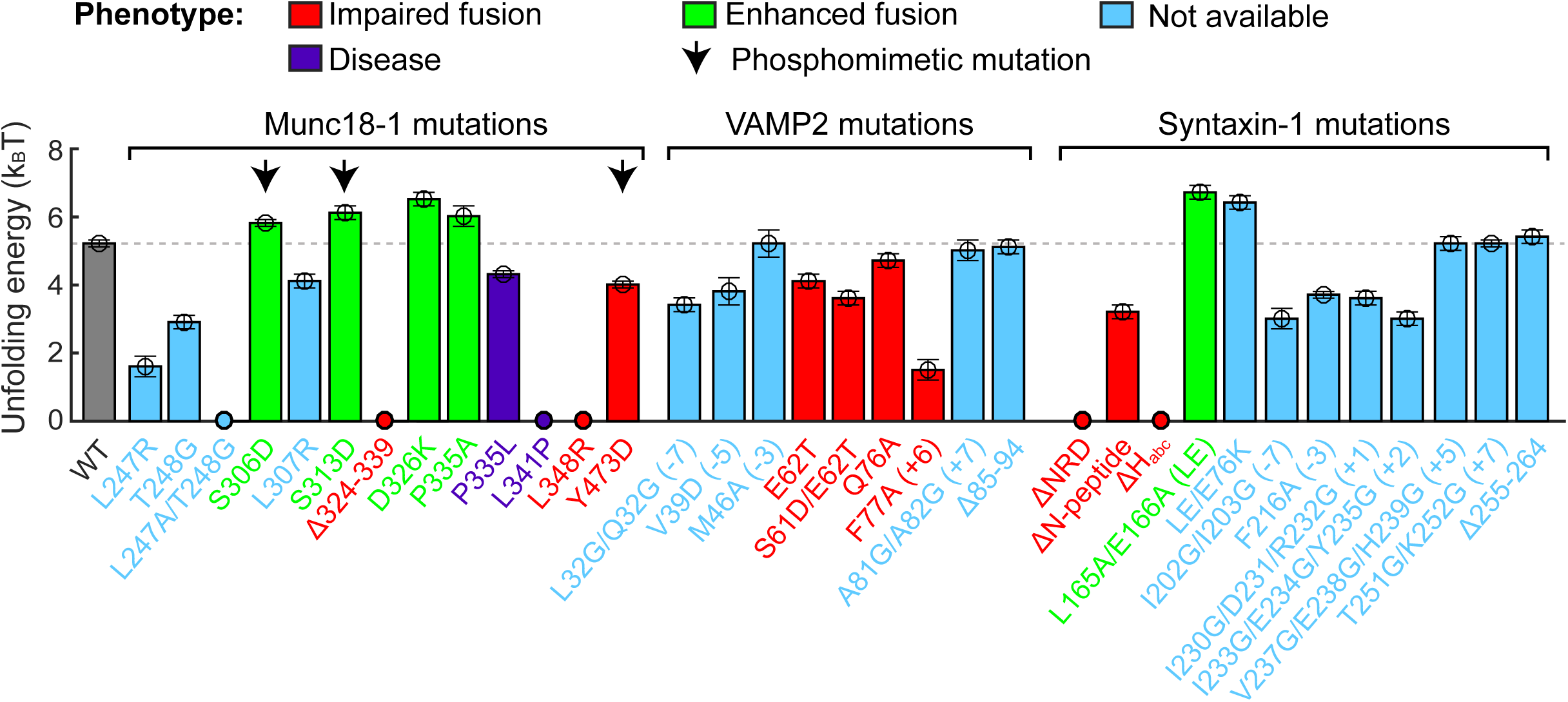
Stability of the template complex correlates with SNARE-mediated membrane fusion and neurotransmitter release. The unfolding energy is derived from the work required to reversibly unfold the template complex (Rebane et al., 2016). The work is measured as the equilibrium force multiplied by the extension change associated with the template complex transition. Numbers in parentheses after SNARE mutant names indicate the layer numbers associated with the corresponding mutations. Error bars indicate standard errors of the mean. See also Table 1 and Figure 3-figure supplement 1.

In the binary Vps33:SNARE crystal structures we reported previously (Baker et al., 2015), only the central regions of each SNARE motif (Qa-SNARE layers −4 to +3; R-SNARE layers −4 to +6) contact the SM template, whereas both ends of each SNARE motif are likely disordered. In the ternary template complex, however, the two SNARE motifs may be correctly zippered all the way to their N-termini. First, −7 layer mutations (VAMP2 L32G/L33G or syntaxin I202G/I203G) destabilized the template complex, as did a −5 layer mutation (VAMP2 V39D) (Figure 3E,I; Figure 4; Table 1). Second, crosslinking the SNAREs at the −6 layer (via syntaxin L205C and VAMP2 Q36C; open arrowhead in Figure 2A; Figure 2-figure supplement 1) (Ma et al., 2015) enhanced the probability of observing the template complex to 0.93 (Figure 2C, #8; Figure 3H). Taken together, these data suggest that the −6 layer is properly aligned in the template complex and that the N-terminal regions from layers −7 to −5 – which are unlikely to contact Munc18-1 but nevertheless contribute to the stability of the complex – are correctly zippered (Figure 2A). By contrast, altering C-terminal regions of the SNARE motifs (VAMP2 A81G/A82G or Δ85-94; syntaxin V237G/E238G/H239G, T251G/K252G, or Δ255-264) did not affect the stability of the template complex (Figure 4 and Table 1). Thus syntaxin regions C-terminal to the +3 layer, and VAMP2 regions C-terminal to the +6 layer, are likely disordered in the template complex.

### Template complex facilitates SNARE assembly

To investigate a potential role for the template complex in SNARE assembly, we repeatedly relaxed and pulled the neuronal Qa-R-SNARE conjugate in the presence of SNAP-25B and, where indicated, Munc18-1. During relaxation, we held the Qa-R-SNARE conjugate at constant mean forces around the equilibrium force of the template complex (Table 1) for up to 60 seconds to afford an opportunity for SNAP-25B binding and SNARE assembly. In the presence of 60 nM SNAP-25B but no Munc18-1, the SNAREs rarely assembled, with a probability of only 0.08 per relaxation (Figure 5A, #1; Figure 5B). Increasing the SNAP-25B concentration to 200 nM increased the assembly probability to 0.41 (Figure 5C, #2-4; Figure 5B). This ‘spontaneous’ (i.e., Munc18-1-independent) SNARE assembly occurred in an all-or-none fashion in the force range of 2-4 pN (Figure 5D, a). Notably, SNAREs misassembled in the absence of Munc18-1 with a probability of ∼0.1, as judged by premature unfolding at low force upon subsequent pulling (Figure 5C, #5; Figure 5B).

**Figure 5.**
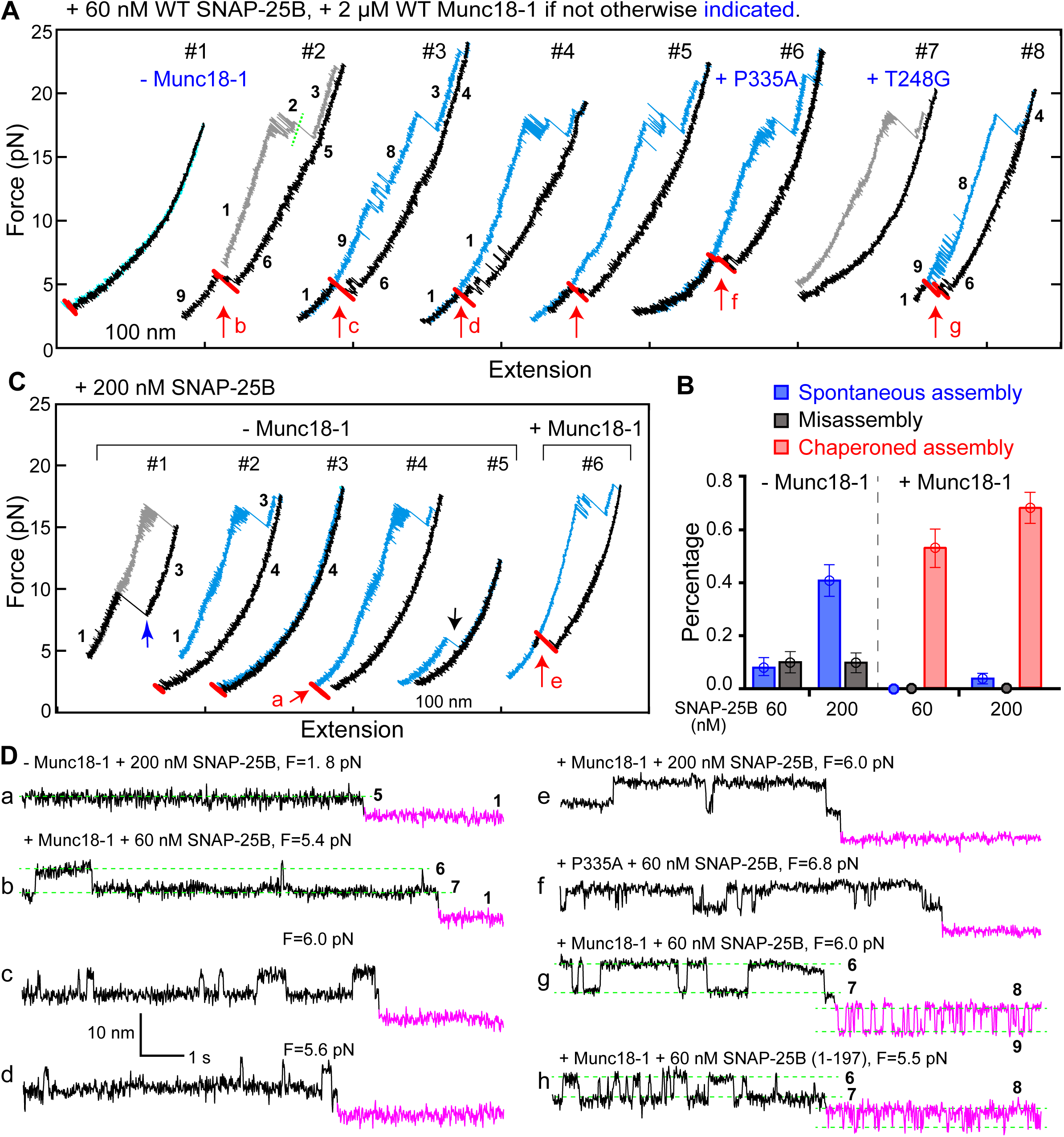
Template complex facilitates snare assembly. (A) Representative FECs obtained in the presence of 60 nM SNAP-25B. Red arrows in Figures 5-7 mark SNARE complex assembly. FECs #2-5 represent consecutive rounds of manipulation of a single Qa-R SNARE conjugate. See also Figure 5-figure supplement 1. (B) Probabilities of Munc18-1-independent (‘spontaneous’) SNARE assembly (blue bars), Munc18-1-chaperoned SNARE assembly (red bars), and SNARE misassembly (black bars). (C) FECs obtained in 200 nM SNAP-25B in the absence or presence of Munc18-1. Arrows mark t-v zippering (blue), disassembly of the misfolded SNARE complex (black), and SNARE reassembly (red). FECs #1-4 are from a single Qa-R SNARE conjugate. (D) Extension-time trajectories at the indicated constant mean forces showing SNARE assembly. All traces were extracted from FEC regions marked with correspondingly labeled red arrows in panels A and B. SNAP-25B-bound states are shown in magenta. See also Figure 5-figure supplement 2.

The addition of 2 μM Munc18-1 increased the frequency of SNARE assembly to 0.53 and 0.68 per relaxation in the presence of 60 nM and 200 nM SNAP-25B, respectively (Figure 5A, #2-#5; Figure 5C, #6). Every SNARE assembly event was preceded by the formation of an intermediate state (Figure 5D, b-e; Video 2; Figure 5B; Table 1). This intermediate had the same average extension relative to the unfolded state, the same equilibrium force, and the same response to mutations as the template complex (Figure 5A, #6-7; Figure 5D, f; Figure 5-figure supplement 1,2). We conclude that in the presence of Munc18-1, the pre-assembled template complex is required for SNAP-25B binding and SNARE assembly.

The template complex greatly accelerated proper SNARE assembly. SNAP-25B bound to the template complex with probabilities of 0.71 and 0.84 per relaxation at 60 nM and 200 nM SNAP-25B, respectively, yielding a binding rate constant of ∼5×10^5^ M^-1^s^-1^. The rate constant is 25-fold greater than that observed in the absence of Munc18-1 (∼2×10^4^ M^-1^s^-1^), presumably because Munc18-1 pre-aligns the N-terminal portions of the syntaxin and VAMP2 SNARE motifs for recognition by SNAP-25B. Consistent with this view, the VAMP2 −7 layer mutations L32G/L33G nearly abolished SNAP-25B binding (Table 1). Notably, we did not observe any misassembly events in the presence of Munc18-1 (Figure 5B). Thus, Munc18-1 enhanced the speed, and probably the accuracy, of SNARE assembly.

### N-terminal regulatory domain of syntaxin stabilizes template complex

Once initiated, the reversible template complex transition (6↔7) typically persisted for over ten minutes at constant mean force, even after the free Munc18-1 in the solution was removed. We suspected that the persistent association between Munc18-1 and the Qa-R-SNARE conjugate was attributable to the NRD of syntaxin, as suggested by previous results (Burkhardt et al., 2008; Shen et al., 2010; Shen et al., 2007; Zhou et al., 2013). Indeed, NRD truncation (ΔNRD) reduced the probability of observing template complex formation from 0.5 to 0.08 (Figure 2C, #5), consistent with the idea that the NRD recruits Munc18-1. The average lifetime of the template complex formed by ΔNRD was also shorter (Figure 3I & Figure 3-figure supplement 3), indicating that the NRD stabilizes the template complex. Unexpectedly, addition of 2 μM NRD in trans was able to rescue the defect: the template complex now formed efficiently (probability = 0.6, N=35) at an equilibrium force close to that of the WT template complex, albeit with slower transition kinetics (Figure 3I). Thus, the NRD can bind to and stabilize the template complex in trans.

Next, we dissected the roles of different NRD regions. Removing the ‘N-peptide’ at the extreme N-terminus of the NRD (Figure 2B) destabilized the template complex (Figures 3I & 4; Table 1), while removing the three-helix bundle Ha_bc_ domain abolished template complex formation altogether (Table 1). By contrast the ‘LE’ mutation (L165A/E166A in the linker region between the Habc domain and the SNARE motif; see Figure 2B) (Dulubova et al., 1999), which promoted SNARE assembly (Table 1) as expected (Burkhardt et al., 2008; Gerber et al., 2008; Ma et al., 2011; Richmond et al., 2001), stabilized the template complex (Figure 3I). Taken together, our results imply that the NRD has a three-fold role in template complex formation (Video 3). When syntaxin is closed (state 6’ in Figure 2B), the NRD inhibits template complex formation (Figure 2-figure supplement 5), as also shown previously (Burkhardt et al., 2008). When syntaxin is partially open (state 6 in Figure 2D), the NRD recruits Munc18-1 for fast folding of the template complex. Finally, once the template complex has formed (state 7 in Figure 2D), the NRD plays a direct stabilizing role. The structural basis for this stabilizing role awaits further investigation.

### Munc18-1 inhibits t-SNARE complex formation

SNARE complex formation in the absence of Munc18-1 was relatively efficient when SNAP-25B was present at high concentrations, as noted above. The addition of Munc18-1, however, reduced the probability of spontaneous SNARE assembly (i.e., assembly not preceded by template complex formation) to 0.04 (Figure 5B). Thus, Munc18-1 not only promotes SNARE assembly via the template complex, but also inhibits spontaneous SNARE assembly. Using a very similar experimental approach, we previously showed that spontaneous SNARE assembly proceeds by a different route (Gao et al., 2012; Zhang et al., 2016a). First, syntaxin binds SNAP-25B to form a t-SNARE complex; then, VAMP2 assembles with the t-SNARE complex in a process called t-v zippering (Gao et al., 2012; Zhang et al., 2016a). In the presence of Munc18-1, t-SNARE complexes were never observed. Thus, Munc18-1 appears to inhibit spontaneous SNARE assembly by suppressing formation of the t-SNARE complex intermediate.

We find that Munc18-1 accelerates SNARE assembly by means of an on-pathway template complex intermediate. A previous model instead proposed that Munc18-1 accelerates t-v zippering (Dawidowski and Cafiso, 2016; Jakhanwal et al., 2017; Shen et al., 2007; Zhang et al., 2016a). To address this possibility directly, we pulled ternary SNARE complexes to generate the t-SNARE complex (state 3) in the presence of 2 μM soluble VAMP2 (Figure 7A). The free VAMP2 molecule rapidly bound the t-SNARE complex (Figure 7B,C), with a binding rate constant (1.6×10^6^ M^-1^s^-1^) close to a previously published value (0.5×10^6^ M^-1^s^-1^) (Pobbati et al., 2006). The binding constant was little changed (2.0×10^6^ M^-1^s^-1^) by the addition of 2 µM Munc18-1. The average t-v zippering force was ∼10 pN with or without Munc18-1 (Figure 5C, #1, blue arrow). Thus, Munc18-1 did not significantly promote t-v zippering.

### Munc18-1 stabilizes the SNARE complex in a partially-zippered state

All of our experiments began with a properly folded SNARE complex (state 1). As noted above, we could drive cycles of disassembly and reassembly by repeatedly pulling and relaxing the Qa-R-SNARE conjugate in the presence of SNAP-25B and Munc18-1. In most cases, the disassembly FECs of the reassembled SNARE complexes were indistinguishable from those of the initial SNARE complexes (Figure 5A, compare #4-5 to #2), suggesting that complete SNARE zippering led to Munc18-1 release (Figure 2D, from state 7 to 1). 14% of the time, however, pulling a reassembled SNARE complex revealed reversible unfolding of the VAMP2 CTD at an unusually low force (4-14 pN, versus 16.5 pN for the initial pull, see blue curves in Figure 5A, #3 & #8). Apparently Munc18-1 was still bound to the assembled SNARE complex (Figure 2D, state 9) (Dulubova et al., 2007); moreover, the unreleased Munc18-1 seemed to stabilize a new partially zippered state (state 8) containing SNAP-25B (Ma et al., 2015). The pulling FEC would therefore be interpreted, in terms of Figure 2D, as 9↔8→3→4.

Occasionally, state 8 was observed at constant mean force (Figure 5A, #8; Figure 5D, g; Video 4). In these cases, reversible template complex formation (6↔7) was followed by reversible CTD folding (8↔9). The 7→8 transition, representing SNAP-25B binding to the template complex, was apparently irreversible. The extension decrease associated with the 7→8 transition was small but significant, implying that the binding of SNAP-25B to the template complex causes additional zippering of syntaxin and VAMP2. The detailed structure of state 8 awaits further investigation.

We predicted that the frequency with which state 8 is observed would be increased by impeding full SNARE zippering and thereby Munc18-1 release. In accord with this prediction, a 9-residue C-terminal truncation of SNAP-25B, which removes layers +7 and +8 of the Qc-SNARE motif and mimics its cleavage by botulinum neurotoxin A (Sutton et al., 1998), increased the probability of detecting state 8 from 0.04 to 0.3 (Figure 5D, h). Along similar lines, we reported recently that the SNAP25B mutations I67N and I67T, both of which alter the +4 layer residue of the Qb-SNARE motif, destabilize SNARE CTD zippering (Rebane et al., 2018). Replacing WT SNAP-25B with either mutant increased the probability of observing state 8 to ∼0.4 (Figure 6B, g). Taken together, our results establish that SNAP-25B binding to the template complex (state 7) leads to the formation of a partially zippered complex containing all three SNAREs and Munc18-1 (state 8), and that the completion of zippering usually expels Munc18-1.

**Figure 6.**
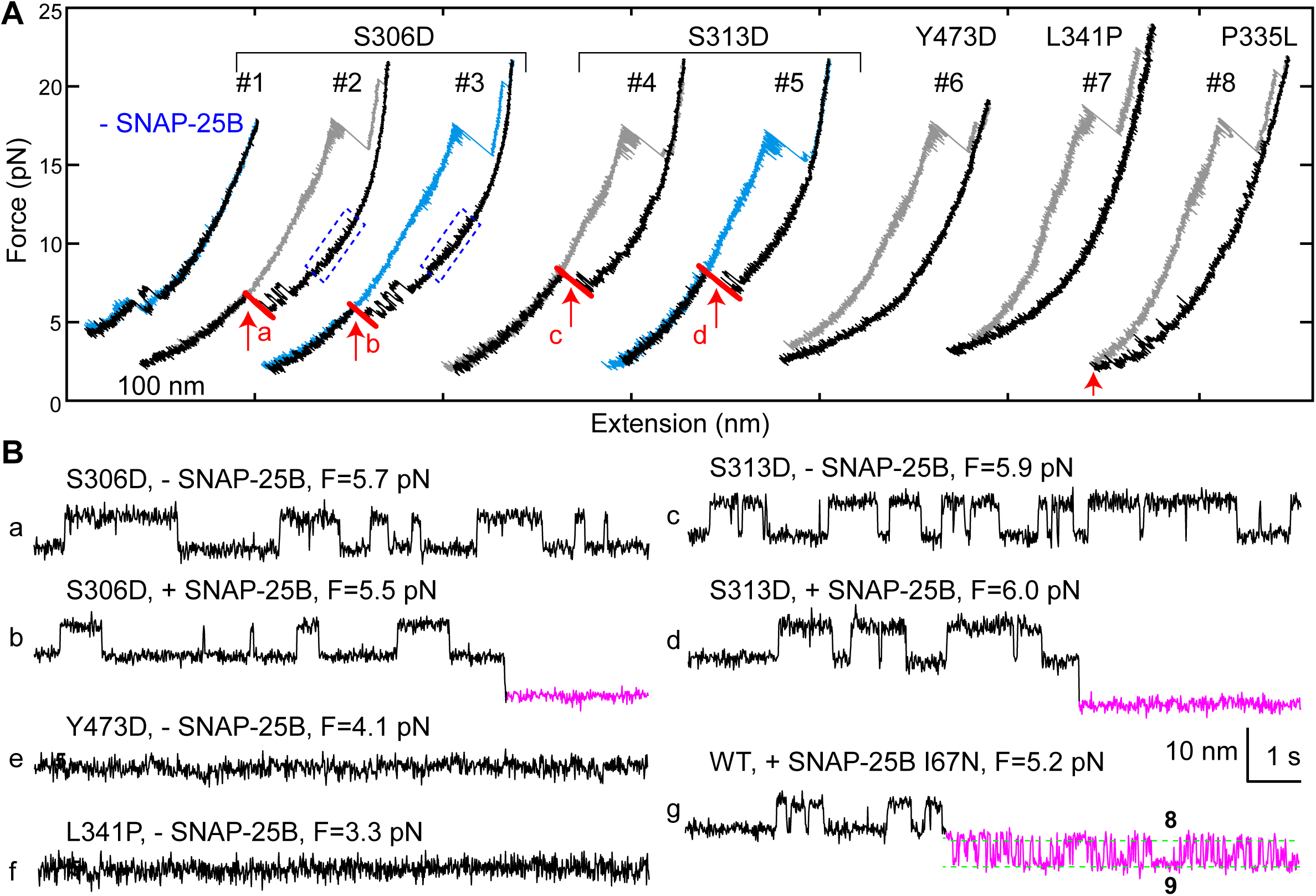
Munc18-1 phosphomimetic and disease mutations altered chaperoned SNARE assembly. (A) FECs for Munc18-1 mutations with 0 nM (#1) or 60 nM (#2-8) SNAP-25B. (B) Extension-time trajectories at the indicated constant mean forces, some of which (a-d) are extracted from panel A. In panels e and f, no template complex formation is observed.

### Function-altering and phosphomimetic mutations

In examining a large number of SNARE and SM mutations (Figure 4 and Table 1), we found that modifications known to compromise SNARE assembly, membrane fusion, and/or neurotransmitter release (Munch et al., 2016; Parisotto et al., 2014; Shen et al., 2007; Walter et al., 2010; Zhou et al., 2013) invariably destabilized the template complex (Figure 4, red bars). Munc18-1 I341P and P335L are, along with SNAP25B I67N and I67P discussed above, four examples of hundreds of Munc18-1 and SNARE mutations associated with epilepsy and other disorders (Stamberger et al., 2016). P335L destabilized, and I341P abrogated altogether, the template complex, leading to impaired Munc18-1-chaperoned SNARE assembly (Figure 4, purple bars; Figure 6A, #7-8; Figure 6B, f). Conversely, mutations known to increase SNARE assembly, membrane fusion, and/or neurotransmitter release (Genc et al., 2014; Gerber et al., 2008; Munch et al., 2016; Parisotto et al., 2014; Sitarska et al., 2017) displayed enhanced template complex stability (Figure 4, green bars). This correlation establishes that the template complex is an important intermediate for membrane fusion in vivo.

The phosphorylation of Munc18 proteins regulates neurotransmitter release and insulin secretion (Genc et al., 2014; Meijer et al., 2018). To explore the mechanism(s) underlying these observations, we examined three phosphomimetic mutations. Phosphorylation of domain 3a residues Ser 306 and Ser 313 (Figure 2A-B) enhances neurotransmitter release and contribute to short-term memory (Genc et al., 2014). Correspondingly, the phosphomimetic mutations S306D and S313D each stabilized the template complex (Figure 4, arrows; Figure 6) and increased the probability of template formation (Table 1). Conversely, the phosphomimetic mutation Y473D, which abrogates membrane fusion in vivo (Meijer et al., 2018), destabilized the template complex (Figures 4 & 6) and reduced the probability of template formation (Table 1). Tyr 473 is located immediately adjacent to the predicted binding pocket for the +6 layer Phe of VAMP2 (Figure 3F) and likely plays a key role in VAMP2 binding. Taken together, these results suggest that Munc18-1 phosphorylation regulates synaptic vesicle fusion by modulating the stability of the template complex.

### Template mechanism is conserved among SM proteins

To generalize our findings, we investigated two other SM proteins, Munc18-3 and Vps33 (Figure 2-figure supplement 1). Munc18-3 and its cognate SNAREs syntaxin 4 (Qa), VAMP2 (R), and SNAP-23 (Qbc) mediate fusion of glucose transporter 4-(GLUT4-) containing vesicles with the plasma membrane, promoting glucose uptake (Bryant and Gould, 2011). Vps33 and its cognate vacuolar SNAREs Vam3 (Qa), Nyv1 (R), Vti1 (Qb), and Vam7 (Qc) mediate membrane fusion in endo-lysosomal trafficking (Wickner, 2010). Like other SNARE complexes (Zorman et al., 2014), the GLUT4 and vacuolar SNARE complexes disassembled stepwise via one or more partially-zippered intermediates (Figure 8A). In the absence of SM proteins, spontaneous SNARE assembly was inefficient, with a probability per relaxation of 0.02 for GLUT4 SNAREs (60 nM SNAP-23) and of 0.04 for vacuolar SNAREs (1 µM Vti1 and 1 μM Vam7) (Figure 8-figure supplement 1). In the presence of 2 µM Munc18-3 or 0.4 µM Vps33, the probability of SNARE assembly increased to 0.44 for Munc18-3 and to 0.65 for Vps33 (Figure 8A, #1-5). Thus, both SM proteins strongly enhance the rate of SNARE assembly. All of the more than 50 Munc18-3-mediated SNARE assembly events we observed in our experiments were mediated by the corresponding template complexes (blue ovals in Figure 8A, #1-3; Figure 8B, a-c). The Munc18-3 template complex displayed a stability (4.3 ± 0.2 kBT) and an extension relative to the unfolded state similar to those of the Munc18-1 template complex. In addition, the Munc18-3 template complex depended on the NRD of syntaxin 4, as NRD truncation reduced the probability of observing the template complex transition to 0.03 (Figure 8B, d). Thus, Munc18-3 and Munc18-1 are quantitatively similar in their ability to chaperone cognate SNARE complex assembly. Importantly, neither Munc18-1 nor Munc18-3 could catalyze the assembly of the other’s cognate SNAREs (Figure 8B, e).

**Figure 7.**
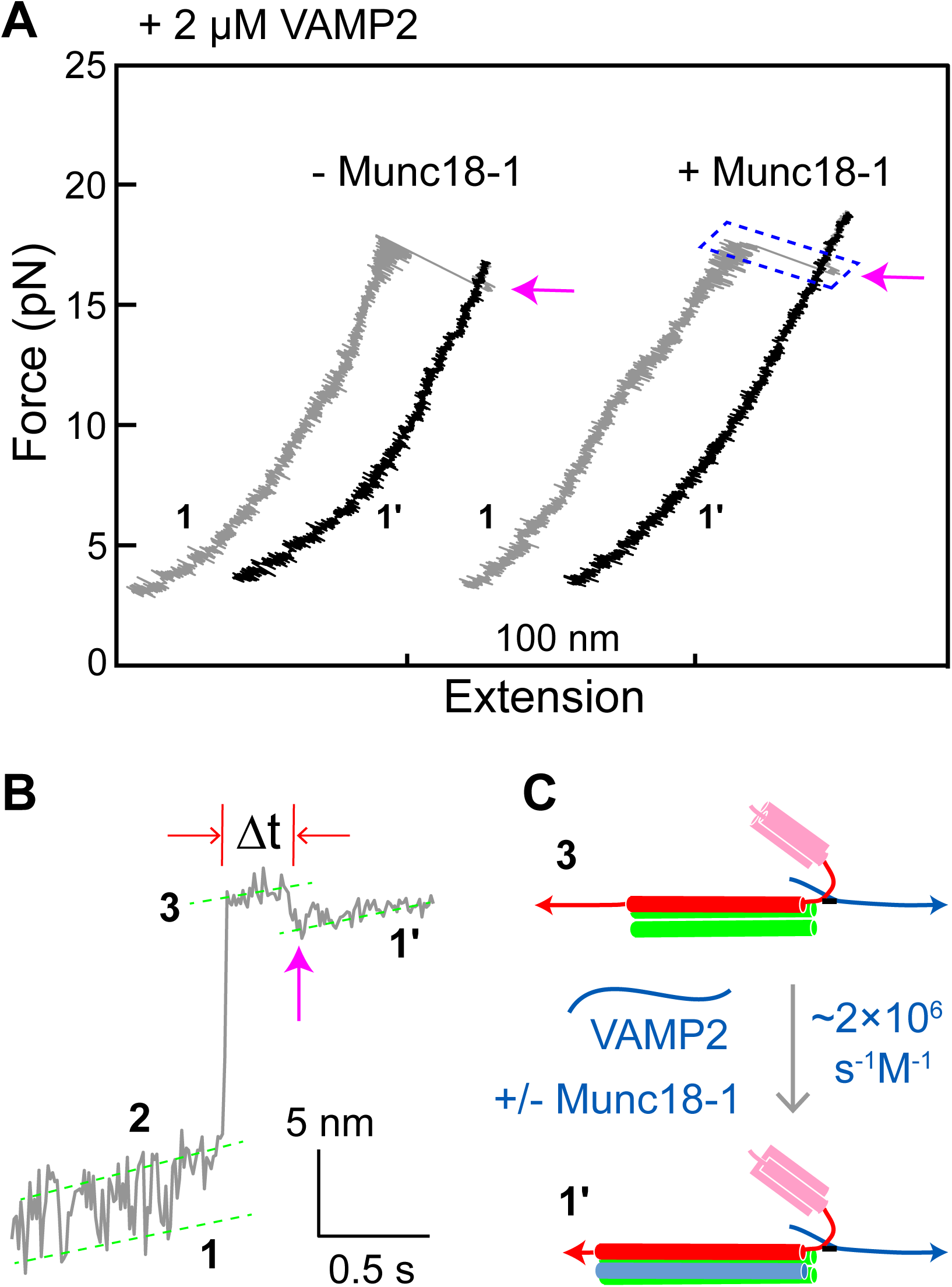
Munc18-1 does not significantly accelerate zippering between t- and v-SNAREs. (A) FECs obtained by pulling single WT SNARE complexes in 2 µM VAMP2 in the solution in the absence or presence of 2 µM Munc18-1. Magenta arrows mark binding of the VAMP2 molecules in the solution to the t-SNARE complexes generated by unzipping the ternary SNARE complex. (B) Close-up view of an extension-time trajectory displaying VAMP2 binding in trans. The trajectory corresponds to the boxed pulling region in A. As observed previously (Ma et al., 2015; Zhang et al., 2016a), VAMP2 binding (indicated by the magenta arrow) induced folding of the disordered C-terminus of the t-SNARE complex, decreasing its extension by 2.3 ± 0.1 nm and generating state 1′ (see C). It took an average time (Δt) of ∼0.3 s for the free VAMP2 in the solution to bind the t-SNARE complex. (C) Diagram illustrating VAMP2 induced t-SNARE folding and extension shortening.

**Figure 8.**
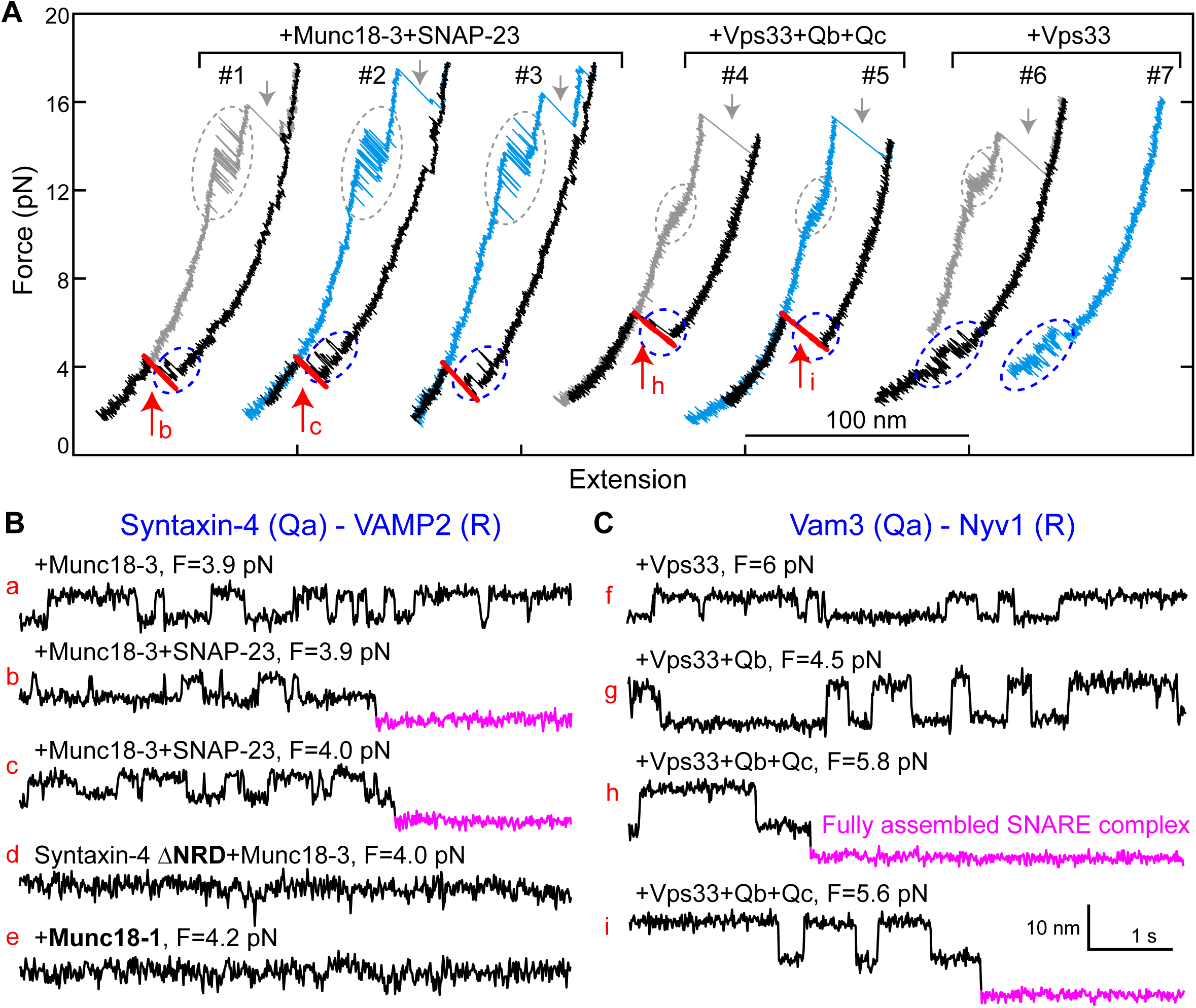
Munc18-3 and Vps33 catalyze SNARE assembly via template complexes. (A) FECs of the Munc18-3 or Vps33 cognate Qa-R SNARE conjugate in the presence of the indicated protein(s). (B) Extension-time trajectories at the indicated constant mean forces, some of which (b, c, h, and i) are extracted from panel A.

We also observed a Vps33-mediated template complex in the absence of Qb-and Qc-SNAREs (blue ovals in Figure 8A, #6-7; Figure 8C, f). Relaxing the Qa-R-SNARE conjugate in the presence of both Vps33 and the Qb-SNARE increased the extension change associated with the template complex transition from 4-6 nm to 7-9 nm (Figure 8C, compare trace g to trace f), indicating that the Qb-SNARE induced further folding of the templated SNAREs. Relaxing the Qa-R-SNARE conjugate in the presence of both the Qb- and Qc-SNAREs triggered assembly of the full SNARE complex from the template complex (Figure 8A, #4-5; Figure 8-figure supplement 2,3). Taken together, these results indicate that Munc18-3 and Vps33 catalyze SNARE assembly by templating SNARE folding and association in a manner analogous to that observed for Munc18-1, in strong support of a conserved templating mechanism underlying SM protein function.

## Discussion

Using geometrically faithful single-molecule experiments, we have mapped out a new pathway for the assembly of neuronal SNARE complexes. As suggested previously (Baker et al., 2015; Sitarska et al., 2017), the key intermediate is a template complex in which the SM protein Munc18-1 serves as the template to arrange the Qa-SNARE syntaxin and the R-SNARE VAMP2 in a Y-shaped conformation with aligned NTDs and splayed CTDs. Although the first 3-4 layers of the NTDs are not expected to interact directly with the template, they nonetheless appear to be properly zippered in the template complex. Our experiments further indicate that the Qbc-SNARE SNAP-25 binds rapidly to the template complex, presumably by recognizing the properly aligned NTDs of the Qa- and R-SNAREs. SNAP-25 binding is occasionally accompanied by a modest amount of further zippering to form a partially-zippered state stabilized by Munc18-1. Finally, full zippering happens in a single, apparently cooperative transition, in most cases accompanied by Munc18-1 release. Like most enzymatic intermediate states, the template complex is relatively unstable (see Materials and Methods for further analysis), preventing it from functioning as a kinetic trap. Nevertheless, our experiments show that it is an obligatory and productive intermediate, which promotes both the speed and the accuracy of SNARE assembly. In addition, the extensive SM-SNARE interactions within the template complex presumably help to prevent the formation of non-cognate SNARE complexes.

Our data appear to be inconsistent with an alternative model in which Munc18-1 binds to and activates the t-SNARE complex to promote t-v zippering (Dawidowski and Cafiso, 2016; Jakhanwal et al., 2017) (Figure 1, upper pathway). We found that Munc18-1 inhibited t-SNARE complex formation and minimally affected t-v zippering, consistent with previous reports (Ma et al., 2013; Pobbati et al., 2006; Zhang et al., 2015). Shen et al. found that fusion between t-liposomes (containing syntaxin:SNAP-25 complexes) and v-liposomes (containing VAMP2) was only stimulated by Munc18-1 after all three were preincubated at 4°C for 3 hours (Shen et al., 2007). This preincubation, under conditions that prevent fusion, was presumably needed to allow formation of the Munc18-1-stabilized partially-zippered SNARE complexes we observe (Figure 1, state v). Preincubating Munc18-1 with t-liposomes alone resulted in little stimulation, inconsistent with the formation of an activated Munc18-1:t-SNARE complex. Finally, t-SNARE complexes, because they are vulnerable to the ubiquitous SNARE disassembly machinery NSF/SNAP (Lai et al., 2017), do not appear to represent plausible intermediates in physiological SNARE assembly pathways.

Our results identify a new role for the NRD of syntaxin: stabilizing the template complex. The stabilizing effect of the NRD is partitioned between its N-peptide and its Habc domain (Figure 2A,B; Figure 4). The stabilizing effect of the N-peptide, which binds to a distal site on Munc18-1 (Burkhardt et al., 2008), is unsurprising, as the N-peptide has long been thought to promote interactions between Munc18-1 and partially or completely folded SNARE complexes (Dulubova et al., 2007; Ma et al., 2015; Shen et al., 2007). However, the stabilizing effect of the Habc domain, even when it is added in trans, is unexpected. This role adds to the others that have been ascribed to the syntaxin NRD and that have complicated efforts to elucidate the physiological neuronal SNARE assembly pathway (Meijer et al., 2012; Shen et al., 2010; Zhou et al., 2013). By contrast, the Qa-SNARE Vam3 does not adopt a closed conformation (Dulubova et al., 2001), a simplifying feature that prompted us to omit its NRD from both our earlier crystallographic studies (Baker et al., 2015) and from the single-molecule experiments reported here. Fortunately the Vps33 template complex was observable in the absence of the Qa-SNARE NRD (Figure 8). Thus, whereas template complexes appear to be a general feature of SM-mediated SNARE assembly, their stabilization via Qa-SNARE NRDs may represent a more specialized elaboration.

Other factors involved in neurotransmitter release may impinge upon the intermediate states we have identified. For example, Munc13-1 plays important roles in opening syntaxin and promoting proper SNARE complex assembly (Lai et al., 2017; Ma et al., 2011; Yang et al., 2015). The opener function of Munc13-1 was circumvented in our studies by two orthogonal strategies, each of which precludes full syntaxin closure. Notably, however, fully closed syntaxin was only marginally more stable than the template complex (7.2 ± 0.2 kBT vs 5.2 ± 0.2 kBT; Figure 2-figure supplement 4). Given that Munc13-1 binds weakly to both syntaxin and VAMP2 at sites likely complementary to those involved in Munc18-1 binding (Lai et al., 2017; Sitarska et al., 2017; Wang et al., 2017), it is attractive to hypothesize that Munc13-1 exerts both its syntaxin opening and SNARE proofreading activities by binding to and stabilizing the template complex (Figure 1, from state i to state iv). Additional factors including complexin and synaptotagmin likely capture the SNARE complex downstream of the template complex, for example by binding to the partially zippered SNARE complex (state v), thereby imposing further regulatory constraints – especially calcium triggering – on synaptic vesicle fusion (Brunger et al., 2018).

A striking finding in this study is the concordance between the effect of mutations, including phosphomimetic mutations, on neurotransmitter release and on the stability of the template complex (Figure 4; Table 1). This concordance strongly supports the hypothesis that the template complex is a physiologically relevant intermediate in SNARE assembly. It is perhaps surprising that many mutations with strong effects on neurotransmitter release have seemingly modest effects on the stability of the template complex. This is, however, readily explained by the exponential dependence of the overall SNARE assembly rate on the stability of the rate-limiting intermediate, by the requirement for multiple SNARE complexes to mediate efficient membrane fusion (Bao et al., 2018; Mohrmann et al., 2010), and/or by the effects of other factors such as Munc13-1 on the stability of the template complex.

The finding that several SM proteins – Munc18-1, Munc18-3, and Vps33 – all catalyze SNARE assembly via a template complex confirms that this is a key conserved function of SM proteins. SNARE zippering, because it involves the coupled folding and assembly of four intrinsically disordered SNARE motifs, is inefficient (Brunger, 2005; Lai et al., 2017). SM proteins, by increasing both the rate and fidelity of SNARE assembly, are likely to be key factors for the control of membrane fusion in vivo. Templated assembly may also resist the disassembly activity of NSF/SNAP.

We propose a working model, using neuronal exocytosis as an example, that places our results in the context of the full fusion machinery (Figure 1). First, SNAREs and SM proteins are recruited to, and thereby concentrated at, the future site of membrane fusion during vesicle docking (Figure 1, state i). Munc13-1 helps bridge vesicle and plasma membranes and recruit SNAREs, and catalyzes opening of the closed syntaxin (state ii) (Lai et al., 2017; Ma et al., 2011; Ma et al., 2013). Subsequently, Munc18-1 binds to the R-SNARE to form the template complex (iv), which may be further stabilized by Munc13-1. Binding of SNAP-25 generates a partially-zippered SNARE complex stabilized by Munc18-1 (state v). Synaptotagmin and complexin likely associate with the partially-zippered SNARE complex, stabilizing it in a primed trans-SNARE complex in preparation for calcium-triggered exocytosis (Sudhof and Rothman, 2009). Finally, calcium triggers fast CTD zippering and Munc18-1 displacement (state vi), inducing membrane fusion.

## Materials and Methods

### SNARE constructs

The cytoplasmic domains of rat neuronal SNAREs, and the SNARE motifs of *C. thermophilum* vacuolar SNAREs, were used. Their sequences are listed below and their domains and crosslinking sites are shown in Figure 2-figure supplement 1. In the sequences below, numbers in parenthesis after each construct name indicate the amino acid numbering in the original protein sequence if there is any truncation, followed by the mutated amino acids, if any, which are also colored red in the sequence. The amino acids in the zero layer are colored cyan. Extra sequences, including linker sequences, are underlined, with Avi-tags or cysteine residues used for crosslinking shown in bold.

### VAMP2 (1-96, N29C)

MSATAATVPPAAPAGEGGPPAPPPNLTS**C**RRLQQTQAQVDEVVDIMRVNVDKVLERDQKLSELDDRADALQAGASQFETSAAKLKRKYWWKNLKMM<u>GGSGNGSGGL</u>**<u>C</u>**<u>TPSRGGD</u>

<u>YKDDDDK</u>

### Syntaxin-1A (1-265, R198C, C145S)

MKDRTQELRTAKDSDDDDDVTVTVDRDRFMDEFFEQVEEIRGFIDKIAENVEEVKRKHSAILASPNPDEKTKEELEELMSDIKKTANKVRSKLKSIEQSIEQEEGLNRSSADLRIRKTQHSTLSRKFVEVMSEYNATQSDYRER**S**KGRIQRQLEITGRTTTSEELEDMLESGNPAIFASGIIMDSSISKQALSEIET**C**HSEIIKLENSIRELHDMFMDMAMLVESQGEMIDRIEYNVEHAVDYVERAVSDTKKAVKYQSKARRKK<u>GGSGNGGSGS</u>**<u>GLNDIFEAQKIEWHE</u>**

### Syntaxin-1A (1-265, I187C, C145S)

MKDRTQELRTAKDSDDDDDVTVTVDRDRFMDEFFEQVEEIRGFIDKIAENVEEVKRKHSAILASPNPDEKTKEELEELMSDIKKTANKVRSKLKSIEQSIEQEEGLNRSSADLRIRKTQHSTLSRKFVEVMSEYNATQSDYRER**S**KGRIQRQLEITGRTTTSEELEDMLESGNPAIFASGIIMDSS**C**SKQALSEIETRHSEIIKLENSIRELHDMFMDMAMLVESQGEMIDRIEYNVEHAVDYVERAVSDTKKAVKYQSKARRKK<u>GGSGNGGSGS</u>**<u>GLNDIFEAQKIEWHE</u>**

### Syntaxin-1A, ΔNRD (187-265, R198C)

ISKQALSEIET**C**HSEIIKLENSIRELHDMFMDMAMLVESQGEMIDRIEYNVEHAVDYVERAVSDTKKAVKYQSKARRKK<u>GGSGNGGSGS</u>**<u>GLNDIFEAQKIEWHE</u>**

### Syntaxin-1A, ΔHabc (Δ27-146, R198C)

MKDRTQELRTAKDSDDDDDVTVTVDR<u>TS</u>GRIQRQLEITGRTTTSEELEDMLESGNPAIFASGIIMDSSISKQALSEIET**C**HSEIIKLENSIRELHDMFMDMAMLVESQGEMIDRIEYNVEHAVDYVERAVSDTKKAVKYQSKARRKK<u>GGSGNGGSGS</u>**<u>GLNDIFEAQKIEWHE</u>**

### SNAP-25B (C85S, C88S, C90S, C92S)

MAEDADMRNELEEMQRRADQLADESLESTRRMLQLVEESKDAGIRTLVMLDE**Q**GEQLERIEEGMDQINKDMKEAEKNLTDLGKF*S*GL*S*V*S*P*S*NKLKSSDAYKKAWGNNQDGVVASQPARVVDEREQMAISGGFIRRVTNDARENEMDENLEQVSGIIGNLRHMALDMGNEIDT**Q**NRQIDRIMEKADSNKTRIDEANQRATKMLGSG

### Syntaxin-4 (1-273, Q194C)

MRDRTHELRQGDNISDDEDEVRVALVVHSGAARLSSPDDEFFQKVQTIRQTMAKLESKVRELEKQQVTILATPLPEESMKQGLQNLREEIKQLGREVRAQLKAIEPQKEEADENYNSVNTRMKKTQHGVLSQQFVELINKSNSMQSEYREKNVERIRRQLKITNAGMVSDEELEQMLDSGQSEVFVSNILKDT**C**VTRQALNEISARHSEIQQLERSIRELHEIFTFLATEVEM**Q**GEMINRIEKNILSSADYVERGQEHVKIALENQKKARKKK<u>GGSGNGGSGS</u>**<u>GLNDIFEAQKIE</u>**

WHE

### Syntaxin-4, ΔNRD (191-273, R206C)

<u>G</u>KDTQVTRQALNEISA**C**HSEIQQLERSIRELHEIFTFLATEVEM**Q**GEMINRIEKNILSSADYVERGQEHVKIALENQKKARKKK<u>GGSGNGGSGS</u>**<u>GLNDIFEAQKIEWHE</u>**

### SNAP-23 (C79S, C80S, C83S, C85S, C87S)

MDDLSPEEIQLRAHQVTDESLESTRRILGLAIESQDAGIKTITMLDEQGEQLNRIEEGMDQINKDMREAEKTLTELNKSSGLSVSPSNRTKNFESGKNYKATWGDGGDSSPSNVVSKQPSRITNGQPQQTTGAASGGYIKRITNDAREDEMEENLTQVGSILGNLKNMALDMGNEIDAQNQQIQKITEKADTNKNRIDIANTRAKKLIDS

### Nyv1 (148-218)

<u>GSS</u>**<u>C</u>**<u>GGG</u>VENNGGDSINSVQREIEDVRGIMSRNIEGLLERGERIDLLVDKTDRLGGSAREFRLRSRGLKRKMWWKNVK<u>GGSGNGSGGG</u>**<u>C</u>**<u>KAAA</u>

### Vam3 (181-252)

<u>GSS</u>**<u>C</u>**<u>GGG</u>LILEREEEIRNIEQGVSDLNVLFQQVAQLVAEQGEVLDTIERNVEAVGDDTRGADRELRAAARYQKRARSRM<u>GGSGNGSGLKNSGGSGSGGNRGGSDSGGSG</u>**<u>GLNDIFEA</u>**

QKIEWHEAAA

### Vti1 (126-190)

<u>GS</u>MLDRSTQRLKASQALAAETEAIGASMLAQLQQQREVIANTTRILYESEGYVDRSIKSLKGIARRM

### Vam7(308-371)

<u>GSQ</u>KLDEQEEYVKDIGVHVRRLRHLGTEIYNAIEQSKDDLDTLDQGLTRLGNGLDKAKALEKKVSGR

### DNA handle preparation

The DNA handle used in our single-molecule experiments is 2,260 bp in length and contains a thiol group (-SH) at one end and two digoxigenin moieties at the other end. The DNA handle was generated by PCR and purified using a PCR purification kit (Qiagen). Both labels were added to the 5′ ends of the PCR primers during synthesis.

### Protein purification

The coding sequences for rat or human syntaxin-1A, VAMP2, Munc18-1, and syntaxin-4 were cloned into pET-SUMO (Invitrogen), which introduced a His6-SUMO tag at the N-termini of the proteins. The coding sequences for rat SNAP-25B and SNAP-23 were cloned into pET-15b (Novagen), which introduced a His6 tag at the N-terminus of the protein. The coding sequence for rat Munc18-3 was cloned into pET-15a (Novagen) and codon-optimized for protein expression in bacteria (Morey et al., 2017). The plasmids were transformed into *Escherichia coli* BL21 (DE3) cells (Agilent Technologies), which were then grown in LB media supplemented with the appropriate antibiotics at 37°C until the OD at 600 nm reached 0.6-0.8. The cells were induced with 1 mM IPTG at 37°C for 5 h. Variants of syntaxin-1A, VAMP2, SNAP-25B and Munc18-1 were prepared using standard PCR-based site-directed mutagenesis (Qiagen).

The neuronal SNARE proteins and Munc18-1 were purified using His-tag affinity purification, as previously described (Gao et al., 2012; Ma et al., 2015). Briefly, the cells were disrupted in HEPES buffer (25 mM HEPES, 400 mM KCl, 10% glycerol, 0.5 mM TCEP, pH 7.7) containing 10 mM imidazole and one tablet of EDTA free protease inhibitor cocktail (cOmplete™, Roche). Cell lysates were cleared by ultracentrifugation. The resulting supernatant was mixed with Ni-NTA resin overnight, after which the resin was washed successively with HEPES buffer containing 20, 40, and 60 mM imidazole. SNAP-25B, VAMP2 and Munc18-1 were eluted in HEPES buffer containing 300 mM imidazole. Syntaxin-1A was eluted in biotinylation buffer (25 mM HEPES, 200 mM potassium glutamate, 300 mM imidazole, pH 7.7) for future biotinylation (see below). For VAMP2 and Munc18-1, the His6-SUMO tags were cleaved by SUMO proteases at 4°C overnight. The cleaved tags were removed by binding to Ni-NTA resin followed by centrifugation.

The *Chaetomium thermophilum* vacuolar SNARE motifs and Vps33 were purified using a previously described protocol with minor modifications (Baker et al., 2015). The *C. thermophilum* SNARE motifs were cloned into a modified pQLinkH vector, resulting in an N-terminal His7-MBP-tag. The plasmids were transformed into *E. coli* C43 (DE3) cells (Lucigen), which were grown in LB media supplemented with ampicillin at 37°C until the OD at 600 nm reached ∼0.6. The cells were induced with 0.5 nm IPTG at 30°C for 4 h and disrupted in lysis buffer (20 mM HEPES, pH 8.0, 350 mM NaCl, 10 mM β-mercaptoethanol and 1 mM PMSF) supplemented with 40 mM imidazole. The lysate was cleared by centrifugation at 17,000 g for 30 min. The His7-MBP-tagged SNARE domains and His7-tagged Vps33 were purified by binding to Ni-NTA resin for several hours, followed by three washes with lysis buffer supplemented with 40 mM imidazole, and elution in lysis buffer supplemented with 300 mM imidazole. For Vps33, the protein was concentrated, followed by size exclusion chromatography on a S200 column equilibrated with gel filtration buffer (20 mM Tris pH 8.0, 250 mM NaCl, 5% glycerol and 0.5 mM TCEP). For the SNARE domains, the His7-MBP tag was removed by incubation with TEV protease with a protein:protease ratio of 20:1 for 3 h at room temperature. The sample was pre-cleared by running on a gravity flow amylose column and concentrated, followed by size exclusion chromatography on a S75 column equilibrated with gel filtration buffer. Residual His7-MBP was removed using a gravity flow amylose column.

After purification, Qa-SNAREs (syntaxin-1A and Vam3) were biotinylated at the Avi-tag in the presence of 50 µg/mL BirA, 50 mM bicine buffer, pH 8.3, 10 mM ATP, 10 mM magnesium acetate, and 50 μM d-biotin (Avidity) at 4 °C overnight (Gao et al., 2012; Jiao et al., 2017).

### SNARE complex formation

To form synaptic SNARE complexes, syntaxin-1A, SNAP-25B, and VAMP2 were mixed in a molar ratio of 0.8:1:1.2 and incubated at 4°C overnight in the HEPES buffer (pH 7.7) with 2 mM TCEP. The SNARE complexes were purified using the His-tag on SNAP-25B (Gao et al., 2012). The quality of the purified neuronal SNARE complex was confirmed by its SDS-resistance in denaturing gel electrophoresis. To form *C. thermophilum* vacuolar SNARE complexes, the His-MBP-tagged Nyv1, Vam3, Vti1 and Vam7, 250 nmol each, were mixed and incubated overnight at 4°C. The complexes were separated from unbound SNAREs by size exclusion chromatography. The His-MBP-tags were cleaved from the SNARE domains using TEV protease and removed by binding to amylose resin. The vacuolar SNARE complex was stored in 20 mM Tris, pH 8.0, 250 mM NaCl, 5% glycerol, 0.5 mM TCEP.

### Crosslinking

We crosslinked the R and Qa-SNAREs at the N-termini of their SNARE motifs and the R-SNARE C-terminus and the 2,260 bp-DNA handle after the SNARE complexes were formed. To this end, both SNARE complexes and DNA handles were treated with 2 mM TCEP for 1 h at room temperature, after which Bio-Spin 6 columns (Bio-Rad) were used to change the buffer to crosslinking buffer A (100 mM phosphate buffer, 500 mM NaCl, pH 5.8) for DNA handles or crosslinking buffer B (100 mM phosphate buffer, 500 mM NaCl, pH 8.5) for SNARE complexes. Next, DNA handles were incubated with 1 mM 2,2’-dithiodipyridine disulfide (DTDP) for 1 h at room temperature to activate the thiol group for the following crosslinking reaction. After incubation, the DNA handle was purified using a PCR purification kit and eluted in crosslinking buffer B to remove excess DTDP. Finally, the SNARE complexes were mixed with the DTDP-treated DNA handles in a 50:1 molar ratio in crosslinking buffer B and incubated at room temperature overnight, as previously described (Gao et al., 2012).

### Single-molecule manipulation experiments

All pulling experiments were performed using dual-trap high-resolution optical tweezers as previously described (Gao et al., 2012; Ma et al., 2015). Briefly, an aliquot of the crosslinked protein-DNA mixture containing 10-100 ng DNA was mixed with 10 μL 2.1 μm diameter anti-digoxigenin antibody coated polystyrene beads (Spherotech) and incubated at room temperature for 15 min. Then the anti-digoxigenin coated beads and 2 μL 1.7 μm diameter streptavidin-coated beads (Spherotech) were diluted in 1 mL PBS buffer (137 mM NaCl, 2.7 mM KCl, 8.1 mM Na2HPO4, 1.8 mM KH2PO4, pH 7.4). Subsequently, the bead solutions were separately injected into the top and bottom channels of a homemade microfluidic chamber as described below. The central channel contained PBS buffer with an oxygen scavenging system comprising 400 mg/mL glucose (Sigma-Aldrich), 0.02 unit/mL glucose oxidase (Sigma-Aldrich), and 0.06 unit/mL catalase (Sigma-Aldrich). A single anti-digoxigenin-coated bead was trapped and brought close to a single streptavidin-coated bead held in another optical trap to form a single SNARE-DNA tether between the two beads.

A single SNARE protein (Qa), SNARE conjugate (Qa-R), SNARE complex, or SNARE/SM complex (collectively called the protein complex below) was pulled or relaxed by moving one of the optical traps at a speed of 10 nm/s. In a typical single-molecule manipulation experiment, a single protein complex was first pulled to a high force to completely disassemble the complex, yielding information on the stability and structure of the complex. Then the complex was relaxed to observe its possible refolding or re-assembly. To better observe the assembly of the template complex or the SNARE complex, during relaxation the protein complex was often held at constant trap separations in a force range of 2-8 pN for various times. The formation probability of the complex generally increased as the waiting time increased. Therefore, the formation probability of the template complex or the SNARE four-helix bundle reported in the main text, including Table 1, was determined with a maximum waiting time of one min if no folding was observed.

### Dual-trap high-resolution optical tweezers

The optical tweezers used in our experiments are home-built and described in detail elsewhere (Gao et al., 2012; Ma et al., 2015). Briefly, the tweezers are assembled on an optical table located in an acoustically isolated, temperature- and air-flow-controlled room. A 1064 nm laser beam from a 4 W Nd:YVO4 diode pumped solid state laser (Spectr-Physics, CA) is expanded by a telescope by about 5 fold, and split by a polarizing beam splitter (PBS) into two orthogonally polarized laser beams. The two beams are reflected by two mirrors and combined by another PBS. One of the mirrors is mounted on a nano-positioning stage that can tip/tilt in two axes with high resolution (Mad City Labs, WI). The combined beams are further expanded by about two fold and collimated by another telescope, and focused by a water immersion 60X objective with a numerical aperture of 1.2 (Olympus, PA), forming two optical traps in a central channel of the microfluidic chamber. One of the optical traps can be moved in the sample plane with sub-angstrom resolution via the nano-positioning stage. The flow cell is formed between two coverslips sandwiched by Parafilm cut into three parallel channels. The top and bottom channels are connected to the central channel by glass tubing. The outgoing laser beams are collected and collimated by an identical objective, split again by a PBS, and projected onto two position-sensitive detectors (Pacific Silicon Sensor, CA), which detect displacements of the two beads in optical traps through back-focal-plane interferometry. The optical tweezers are calibrated before each single-molecule experiment by measuring the Brownian motion of the trapped beads, which yields the power-spectrum density distributions of bead displacements. The force constants of optical traps are determined by fitting the measured power-spectrum density distributions with a Lorentzian function.

### Circular dichroism (CD) spectra of Munc18-1

CD spectra of WT and mutant Munc18-1 proteins were measured in 20 mM phosphate buffer using an Applied Photophysics Chirascan equipped with a 2 mm quartz cell. The readings were made at 1 nm intervals, and each data point represents an average of 6 scans at a speed of 120 nm/min over the wavelength range of 190 to 250 nm.

### Derivations of protein unfolding energy and folding and unfolding rates from force-dependent measurements

Our methods of data analysis and the relevant Matlab codes are described in detail elsewhere (Gao et al., 2012; Rebane et al., 2016). Briefly, the extension-time trajectories obtained at constant trap separations or mean forces were first analyzed by two-state hidden-Markov modeling (McKinney et al., 2006; Zhang et al., 2016b), which revealed the idealized state transitions, extension changes, unfolding probabilities, and folding and unfolding rates. These measurements were used to derive the folding intermediates and their associated energy and kinetics.

We quantified the structural change of a single protein based on the measured force and extensions. The control parameter of our pulling experiment is the separation between two optical traps (*D*). Given the trap separation, the extension (*X*) and tension (*F*) of the protein-DNA tether are calculated as

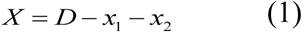

And

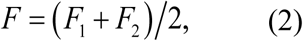

respectively, where*x*1 and*x*2 are displacements of the two beads in optical traps, and *F*1 and *F*2 are the corresponding forces applied to the beads. Both bead displacement *x* and the force *F* are derived from voltage outputs of the position-sensitive detectors after proper calibrations. In Eq. (1), we have defined a default relative trap separation by neglecting the contribution of constant bead diameters. It is this relative trap separation that is shown in Figure 2-figure supplement 2. As a protein *m*olecule unfolds, its extension (*x m*) increases, which leads to retraction of both beads in their optical traps and the accompanying decrease in tension (Figure 2-figure supplement 2). Thus, during protein folding and unfolding transitions, the tether tension changes in an out-of-phase manner with respect to the tether extension, and thus is state-dependent. In the constant trap separation, the mean force is defined as the mean of the average forces associated with the folded and unfolded states (Rebane et al., 2016).

We modeled the unfolded peptide and the DNA handle by a worm-like chain model. Based on this model, the stretching force *F* and the entropic energy *E* of a semi-flexible polymer chain are related to its extension *x*, contour length *L*, and persistence length *P* by the follow formulae

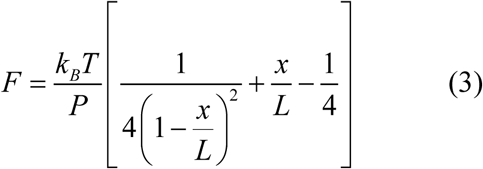

and

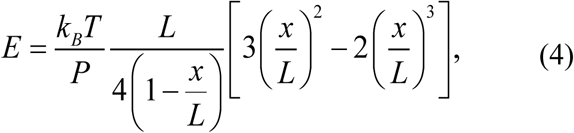

respectively. We adopted a persistence length 40 nm for DNA and 0.6 nm for the unfolded polypeptide (Gao et al., 2012; Ma et al., 2015; Rebane et al., 2016). Because the DNA extension (*x*_*DNA*_) is known given a force or trap separation via Eq. (3), the extension of the protein can be calculated as *x*_*m*_ = *X* – *x*_*DNA*_ The protein extension generally comprises the extensions of the unfolded *p*olypeptide portion (*x _p_*) and the folded portion (*H*) of the protein if any, or *x _m_* = *x p H*. The former can be again calculated by Eq. (3), given the contour length of the unfolded polypeptide (*L*_*p*_), whereas the latter can be treated as a force-independent constant, or a *h*ard core of the protein (Rebane et al., 2016). Here the size of the hard core is determined from the two pulling sites on the folded protein portion, which changes with the protein state. Thus, the contour length of the unfolded polypeptide and the size of the folded protein portion are correlated and can be determined based on a structural model for protein transitions. To derive the structure of the template complex, we assumed a hard core size of 3 nm for the folded template complex, as determined from the structure-based model (Baker et al., 2015). Consequently, we could determine the contour length of the polypeptide chain in the Qa-R conjugate that is either free or bound by Munc18-1. The number of amino acids in a polypeptide is its contour length divided by the contour length per amino acid, which is chosen to be 0.365 nm (Gao et al., 2012; Rebane et al., 2016). The number of amino acids in the completely unfolded SNARE state 5 is known (Figure 2-figure supplement 1), which helps derive the structure of the template complex based on the extension change during the template complex transition. We determined that 87 (±2, S.D.) amino acids are sequestered in the folded template complex, including the N-terminal loop formed between syntaxin-1 and VAMP2 due to crosslinking. Based on our construct design (Figure 2-figure supplement 1), this length is consistent with the structure of the predicted template complex (Figure 1A).

Similarly, we modeled the total free energy of the whole dumb-bell system in optical traps, or

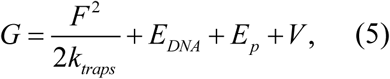

where the first term represents the potential energy of the two beads in optical traps with *k*_*traps*_= *k*_1_*k*_2_ **/** (*k*_1_ + *k*_2_) the effective force constant of the two traps, the second and third terms are entropic energies of the DNA handle and the unfolded polypeptide, respectively, calculated by Eq. (4), and the last term is the free energy of the protein at zero force. Based on the Boltzmann distribution, the protein unfolding energy Δ*V* can be determined by fitting the measured unfolding probability using Eq. (5). Equation (5) can be similarly applied to the transition state of protein folding (Rebane et al., 2016). With Kramers’ rate equation, the folding and unfolding rates are calculated. By fitting the calculated rates to the measured rates, we derive the energy and conformation of the transition state, which also yield the folding and unfolding rates at zero force. Complete data sets from individual molecules are separately fit and the unfolding energies and transition rates, typically averaged over more than three different molecules, are reported (Table 1). The average folding rates and unfolding rates of the WT and mutant template complexes fall in the ranges of 17-568 s^-1^ and 0.1-10 s^-1^, respectively, with a standard error typically close to the corresponding average rate for each template complex.

### Estimation of the affinity between VAMP2 and Munc18-bound syntaxin

The N-terminal crosslinking between Qa-and R-SNAREs used in our assay is crucial for us to observe and characterize the template complex. The crosslinking destabilizes the closed syntaxin, thereby bypassing the requirement for Munc13-1, mitigates SNARE misassembly, for example, formation of various anti-parallel SNARE bundles (Lai et al., 2017), and avoids nonspecific VAMP2-Munc18-1 interactions (Sitarska et al., 2017). Therefore, the crosslinking simplifies our experimental design and data interpretation. Three lines of evidence suggest that the crosslinking does not compromise the major conclusions derived from our assay. First, the stability of the template complex does not depend upon the crosslinking site used in our assay (at R198C or I187C, Figure 2-figure supplement 1). This observation suggests that the crosslinking does not alter the structure of the template complex and is likely located at a disordered region. Second, the derived template model recapitulates many distinct features of the fusion machinery, including its dependence upon NRD, phosphorylation, and various mutations. Finally, the crosslinking increases the local SNARE concentration around Munc18-1, which mimics the environment of SNARE assembly and membrane fusion in vivo due to vesicle tethering and SNARE recruitment. For example, Munc13-1 essentially crosslinks both syntaxin and VAMP2 by simultaneously binding the two (Figure 1).

The effective concentration due to the crosslinking can be quantified, which is used to estimate the binding affinity between VAMP2 and partially-closed syntaxin in the absence of crosslinking (Zhang et al., 2016a). To derive the local concentration of the crosslinked VAMP2 (at R198C) around the partially-closed syntaxin, we made three assumptions: 1) the partially-closed syntaxin has a conformation similar to the conformation seen in the crystal structure of the Vps33:Vam3 complex (Baker et al., 2015); 2) VAMP2 binding kinetics is dominated by insertion of Phe 77 into the F-binding pocket on the Munc18-1 surface, as is supported by our data; and 3) the unfolded SNARE polypeptides are described by a Gaussian chain model. Thus, VAMP2 binding to the partially closed syntaxin can be modeled by Phe 77 binding to the F-binding pocket while Phe 77 is tethered to the −7 layer of Vam3 (i.e., the N-terminus of the main Vam3 helix in the Vps33:Vam3 structure, PDB code 5BUZ) via a polypeptide linker. The length of the linker is 54 amino acids based on our Qa-R SNARE conjugate, or *L*=19.7 nm in terms of the contour length. The distance between the F-binding pocket and the tethering point (*R*) is measured to be 6.34 nm. Therefore, the effective concentration *c* of the tethered Phe 77 around its binding pocket is calculated as *c*=3.7×10^-4^ M, using the following formula

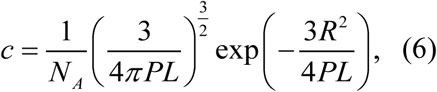

Where *N*_*A*_ is Avogadro’s number and *P*=0.6 nm is the persistence length of the polypeptide. Due to coupled binding and folding, the folding and unfolding rates of the template complex we measure should be equal to the binding and dissociation rates estimated here. Therefore, the folding rate *k*_*f*_=*k*_*on*_×*c*, where *k*_*on*_ is the bimolecular rate constant for VAMP2 binding to the partially-closed syntaxin. Using our measured folding and unfolding rates, we calculated the VAMP2 binding rate constant as *k*_*on*_= 3.5×10^5^ s^-1^M^-1^, energy as 13.1 k_B_T, or a VAMP2 dissociation constant as 2 µM. Supposing that VAMP2 binding to the fully closed syntaxin requires an energy gain of 4.6 k_B_T compared to the partially closed syntaxin (corresponding to the energy difference between the two syntaxin states), the VAMP2 binding affinity to the fully closed syntaxin is estimated to be 200 µM.

## Data and software availability

For custom programs and scripts used in this study, please contact Dr. Yongli Zhang.

## Acknowledgements

We thank J. Rothman and A. Horwich for discussion, G. Shimamura for technical assistance, and D. Fasshauer for providing the plasmid for Munc18-3 purification. This work was supported by NIH grants R01GM093341 and R01GM120193 to Y.Z., T32GM007223 to J.J., and R01GM071574 to F.M.H., and by the German Research Foundation (DFG) grant PO2195/1-1 to S.A.P.

## Author contributions

Conceptualization, J.J., M.H., S.A.P., R.W.B., F.M.H., and Y. Z.; Investigation, J.J., M.H.,

S.A.P., R.W.B., Y.X., H.Q., Y.X., Y.W., H.J.; Writing, J.J., M.H., S.A.P., F.M.H., and Y. Z.;

Software, J.J. and Y.Z.; Formal analysis, J.J., M.H., and Y.Z.; Visualization, J.J., M.H., T.J.E.,

F.M.H., and Y.Z.; Funding Acquisition, S.A.P., F.M.H., and Y.Z.; Supervision, F.M.H., and Y.Z.

**Video 1. SNARE complex unfolding and subsequent template complex formation as inferred from single-molecule measurements.**

The proposed state transitions associated with FEC #2 in Figure 2C or Figure 2-figure supplement 2 are simulated.

**Video 2. Template complex facilitates SNAP-25B binding and SNARE assembly**

The extension at a constant mean force of 6.0 pN corresponding to trace c in Figure 5D and its associated state transition are simulated. For simplicity, only the right bead was simulated to move in response to SNARE conformational changes in Video 2-4. In reality, the left bead moved synchronously with the right bead, but in an opposite direction, as shown in Video 1.

**Video 3. Inferred conformational transition from closed syntaxin to the template complex.**

**Video 4. SNAP-25B binding to the template complex occasionally forms a partially zippered SNARE complex.**

The state transitions associated with the extension trace g in Figure 5D are simulated.

## SUPPLEMENTAL INFORMATION

**Figure 2-figure supplement 1.**
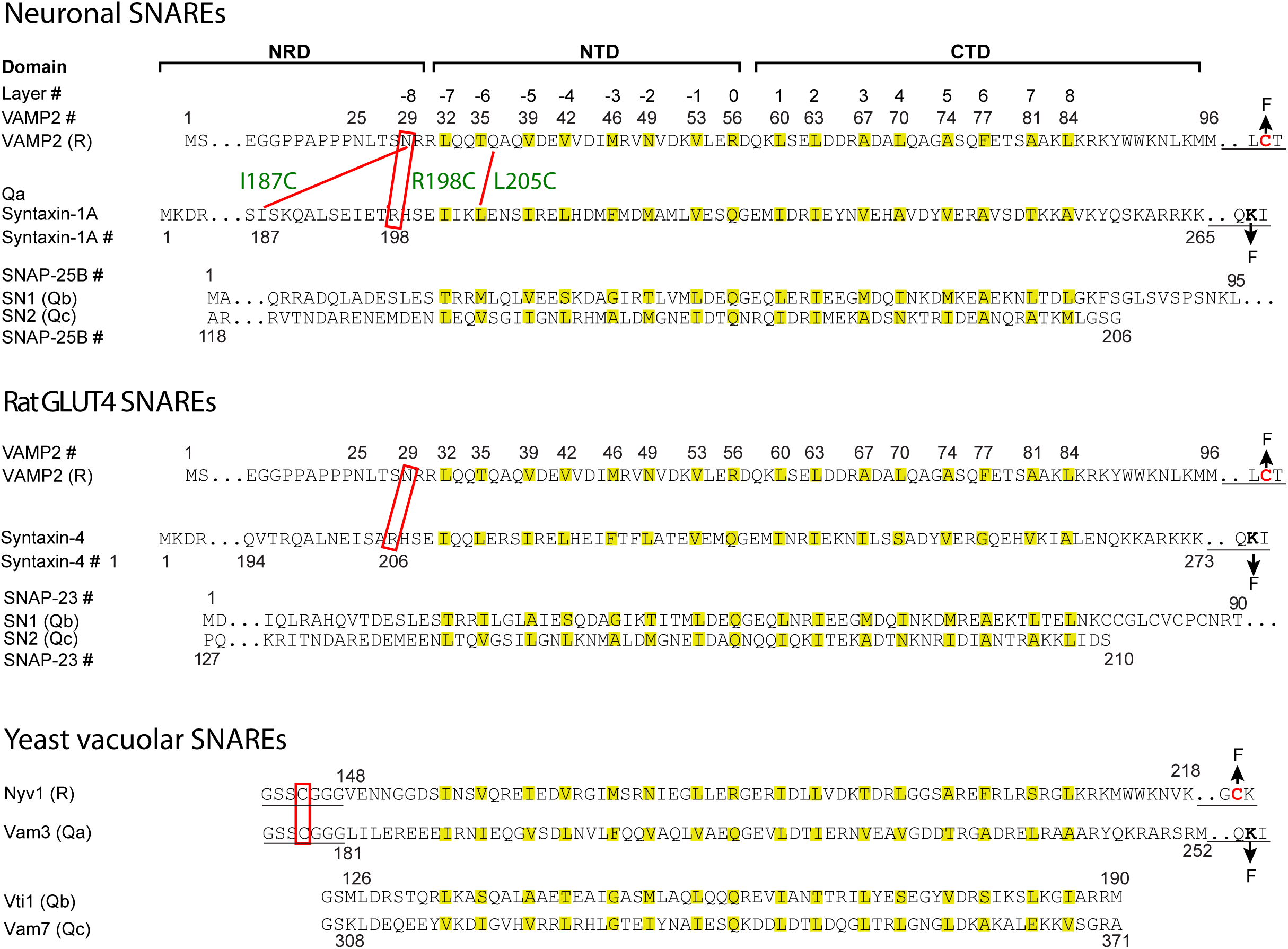
Sequences, domains, and crosslinking sites of the SNARE proteins used in this study. The amino acids in hydrophobic layers (from −7 to +8) and the central ionic layer (0 layer) are colored yellow. The underlined sequences are added to facilitate crosslinking of Qa and R SNAREs, crosslinking of R SNAREs to DNA handles, and attachment of Qa SNAREs to bead surfaces (Figure 2A). The pulling sites are indicated by arrows labeled by F (for Force). The crosslinking sites are indicated by red rectangles or red lines. For neuronal SNAREs, the three crosslinking sites tested are designated by their corresponding syntaxin residues (R198C, I187C, and L205C, Figure 2B,C). N-terminal crosslinking facilitated formation of the template complex by destabilizing the closed syntaxin, increasing the local SNARE concentrations, and minimizing nonspecific Munc18-1-VAMP2 and SNARE-SNARE interactions (Brunger, 2005; Lai et al., 2017; Sitarska et al., 2017). To facilitate assembly of the Qa-R conjugate and the SNARE-DNA tether, we typically crosslinked the Qa- and R-SNAREs in purified SNARE complexes and then removed the Qbc-SNAREs by unfolding the complexes. The force-extension curves (FECs) corresponding to the first pull to unfold these pre-assembled SNARE complexes are shown in gray throughout the text.

**Figure 2-figure supplement 2.**
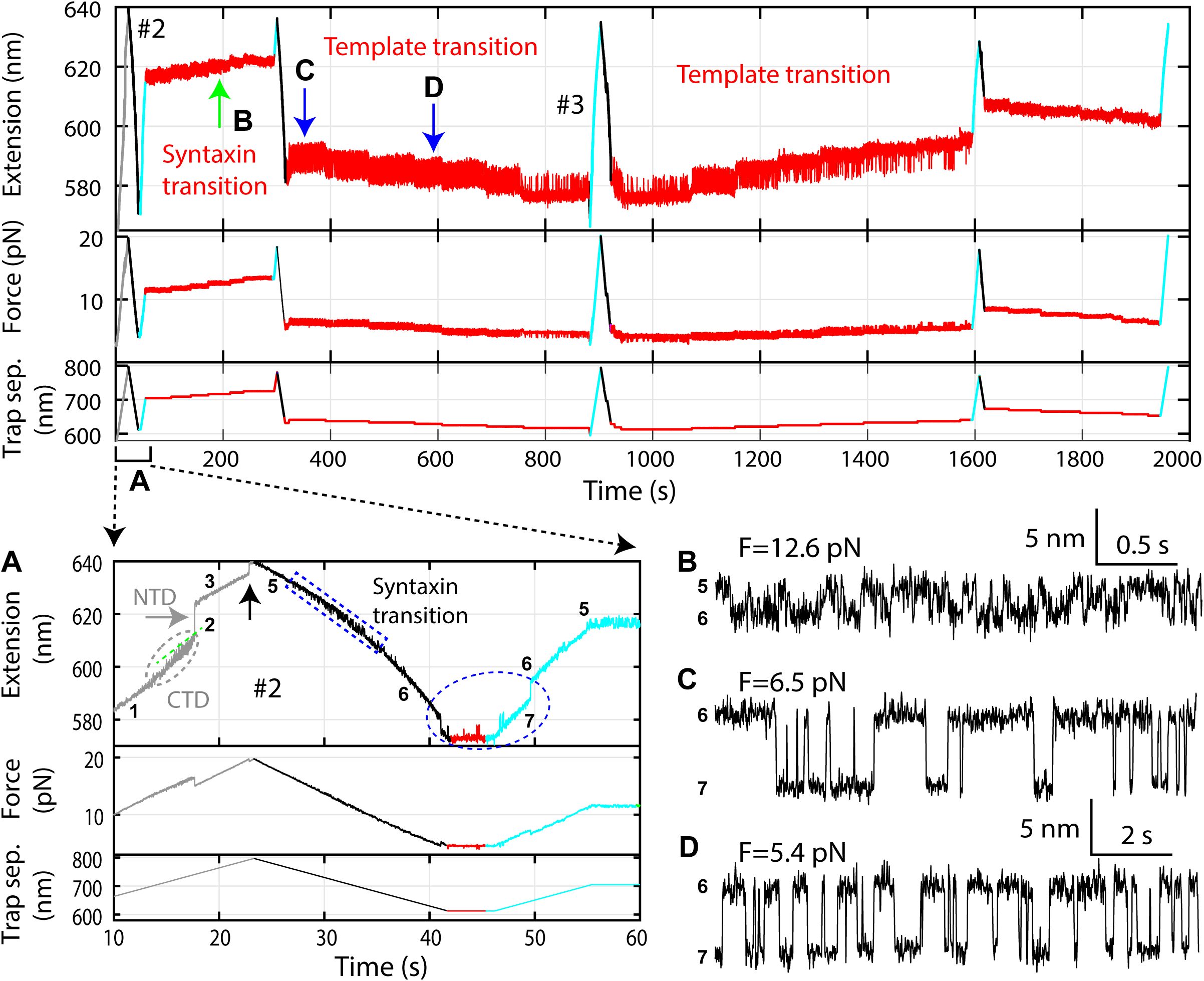
Time-dependent extension (top panel), force (middle panel), and trap separation (bottom panel) for a typical experiment to test template complex formation. Data here and FECs #2 and #3 in Figure 2C are acquired on the same Qa-R SNARE conjugate, with the same pulling round numbering. Close-up views of different time regions indicated by A-D are shown: (A) Close-up view of the first round of pulling and relaxation. Regions of characteristic transitions are indicated: the CTD transition by the gray oval, the NTD unfolding by the gray arrow, the syntaxin unfolding and SNAP-25B dissociation by the black arrow, the syntaxin open-partial closing transition by blue rectangle, and the template complex transition by the blue oval. States associated with different extensions are indicated by the corresponding state numbers (Figure 2D). Note that binding of Munc18-1 to the NRD in state 5 did not cause any change in the extension of the Qa-R conjugate compared to the state 4, thus could not directly be detected by our assay. (B) The extension-time trajectory demonstrates reversible partial closing of syntaxin at a constant mean force of 12.6 pN. (C-D) Extension-time trajectories at two constant trap separations or mean forces showing reversible folding and unfolding of the template complex. Throughout the figures, all FECs and time-dependent trajectories were mean-filtered with a time window of 10 ms, except for the extension-time trajectories with syntaxin opening-partial closing transitions, which were filtered with a time window of 3 ms.

**Figure 2-figure supplement 3.**
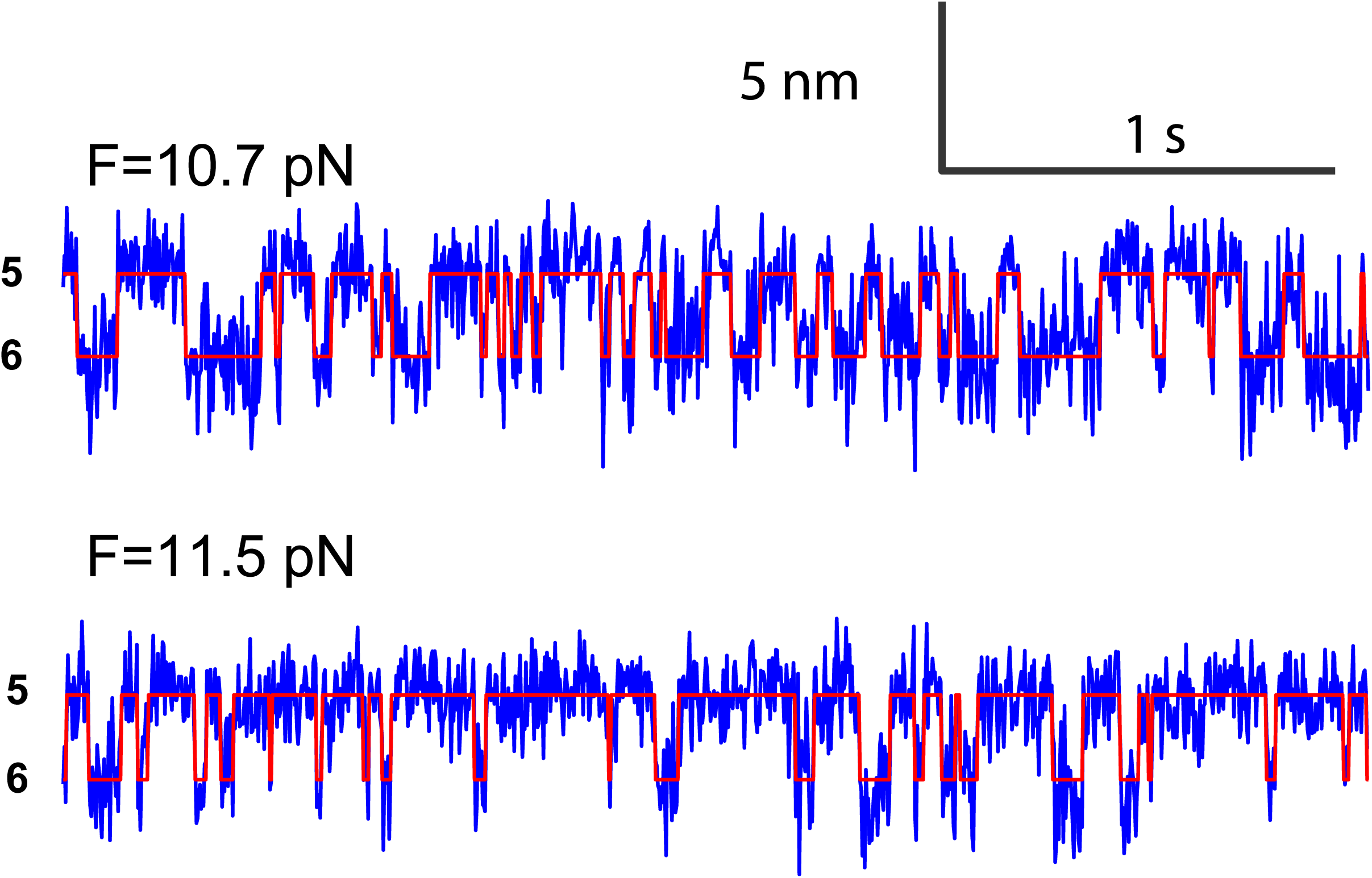
Extension-time trajectories at two constant mean forces (F) showing the opening-closing transition of the partially closed syntaxin molecule.

**Figure 2-figure supplement 4.**
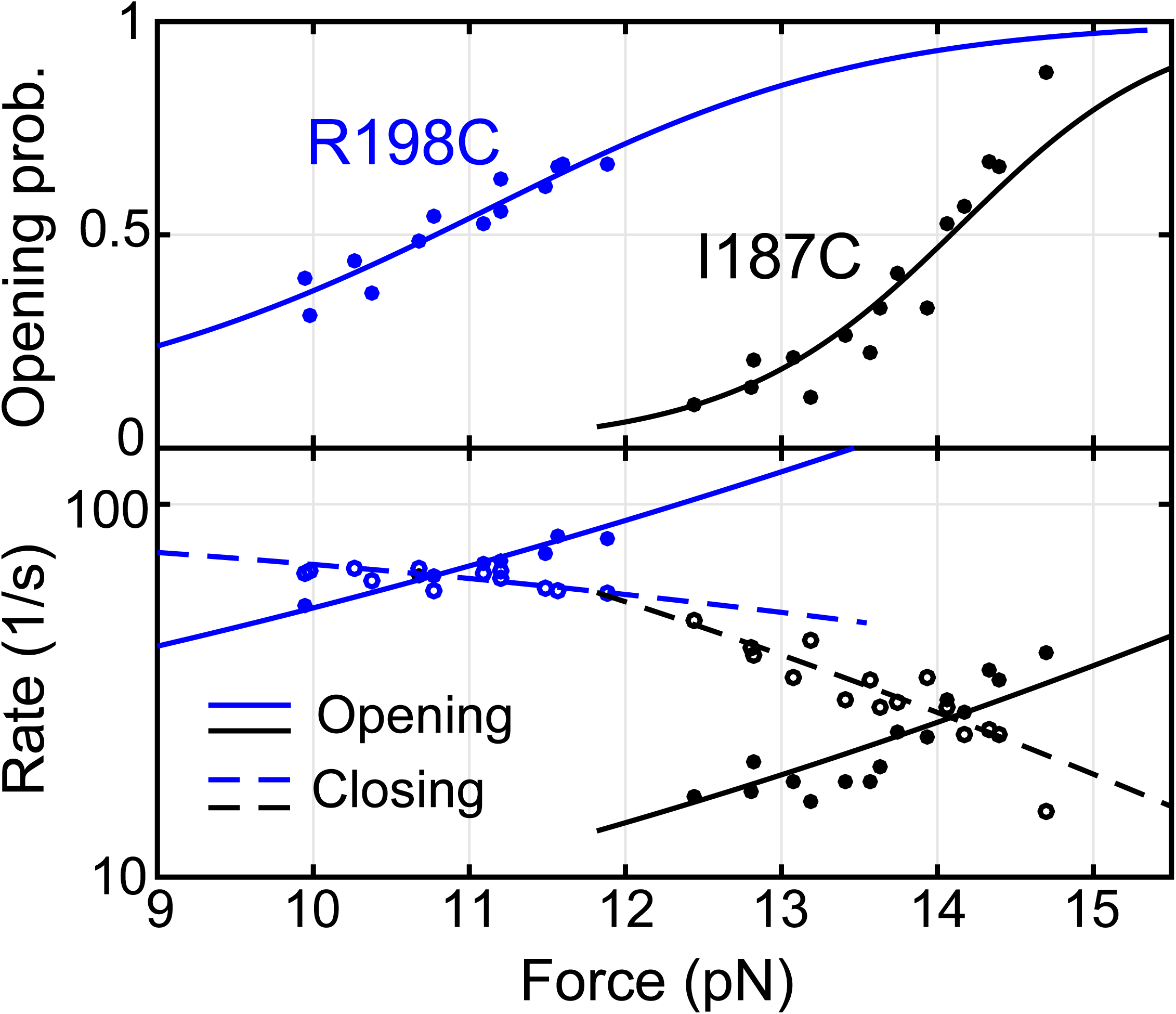
Force-dependent syntaxin opening probabilities (top panel) and opening and closing rates (bottom panel) obtained by pulling syntaxin from the two N-terminal sites, R198C and I187C (Figure 2-figure supplement 1). Curves are best model fits to derive the energies and kinetics at zero force associated with the transitions, with solid curves for unfolding and dashed curves for folding. The unfolding energies of the closed syntaxin (7.2 ± 0.2 kBT) and of the partially closed syntaxin (2.6 ± 0.2 kBT) are much smaller than the dissociation energy between syntaxin and Munc18-1 previously measured based on a two-state binding and unbinding process (∼22 k_B_T) (Burkhardt et al., 2008; Sitarska et al., 2017). Our data revealed an intermediate state for the association and dissociation process, in which syntaxin is open, but Munc18-1 remains bound to syntaxin, likely to the NRD (Ma et al., 2015) (Figure 2D, state 5).

**Figure 2-figure supplement 5.**
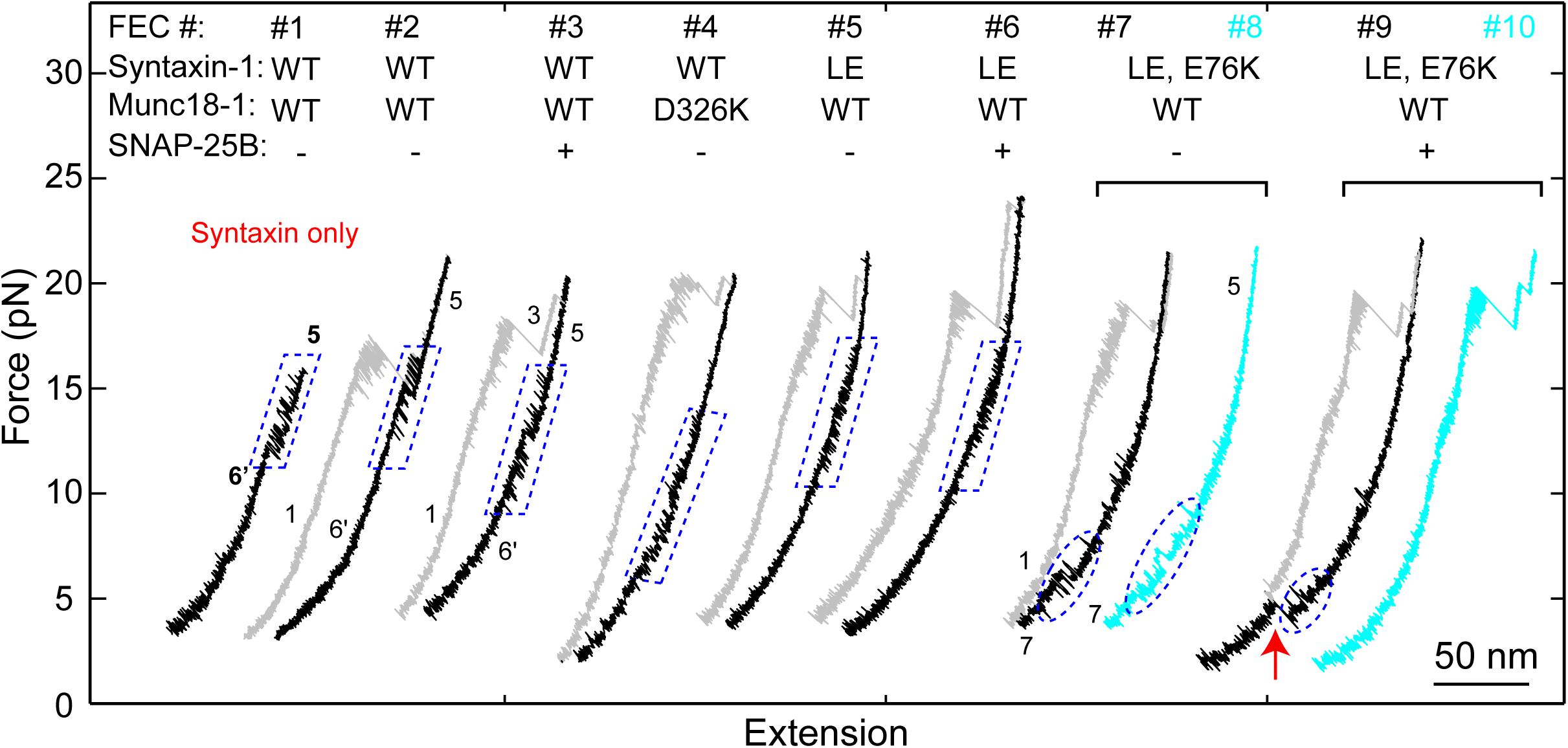
FECs of Qa only (#1) or the Qa-R SNARE conjugate (other FECs) pulled from Site I187C in the absence (-) or presence (+) of 2 μM Munc18-1 or 60 nM SNAP-25B. The wide-type (“WT”) syntaxin-1 here denotes syntaxin-1A (a.a. 1-265, I187C, C145S; see “SNARE protein constructs”), with additional mutations indicated. FECs in each bracket were obtained on the same Qa-R conjugate. Different transitions are marked: blue dashed parallelograms for syntaxin opening-closing transitions, blue dashed ovals for template complex transitions, and the red arrow for SNAP-25B binding and full SNARE assembly. To pull syntaxin alone, we directly crosslinked the DNA handle to syntaxin I187C in the absence of VAMP2. The resultant FEC (#1) revealed the same high-force transition as those obtained by pulling the Qa-R conjugate crosslinked at I187C, indicating that the high-force transition indeed resulted from transition of syntaxin alone (compare to the syntaxin transition in #2-3). The WT syntaxin was fully closed by Munc18-1, which inhibited formation of the template complex (#2-3) in the absence of SNAP-25B (#2) and in the presence of SNAP-25B (#3). Munc18-1 mutation D326K destabilized the closed syntaxin, but was not sufficient to open the closed syntaxin for template complex formation (#4). The LE mutation (L165A/E166A) decreased the extension change associated with the syntaxin transition (compare #5 to #1-3), indicating that the LE mutation destabilized the closed syntaxin conformation (Burkhardt et al., 2008). Nevertheless, neither template complex formation (#5 and #6) nor Munc18-1-chaperoned SNARE assembly (#6) was observed, suggesting that syntaxin with the LE mutation is still largely closed, consistent with recent results (Colbert et al., 2013). Therefore, we made an additional mutation E76K that was expected to further weaken the closed syntaxin conformation (Figure 2B). Indeed, the combined mutation partially opened the syntaxin conformation and promoted formation of the template complex, as seen from small high force transition and appearance of the low force transition, respectively (#7 and #8). Correspondingly, Munc18-1-chaperoned SNARE assembly was observed in the presence of SNAP-25B (red arrow) in a manner that depended on the template complex (#9 and #10). Under this condition, the probabilities of detecting the partially closed syntaxin and the template complex were 0.66 and 0.39, respectively, and the probability of detecting SNARE assembly after template complex formation was 0.48.

**Figure 2-figure supplement 6.**
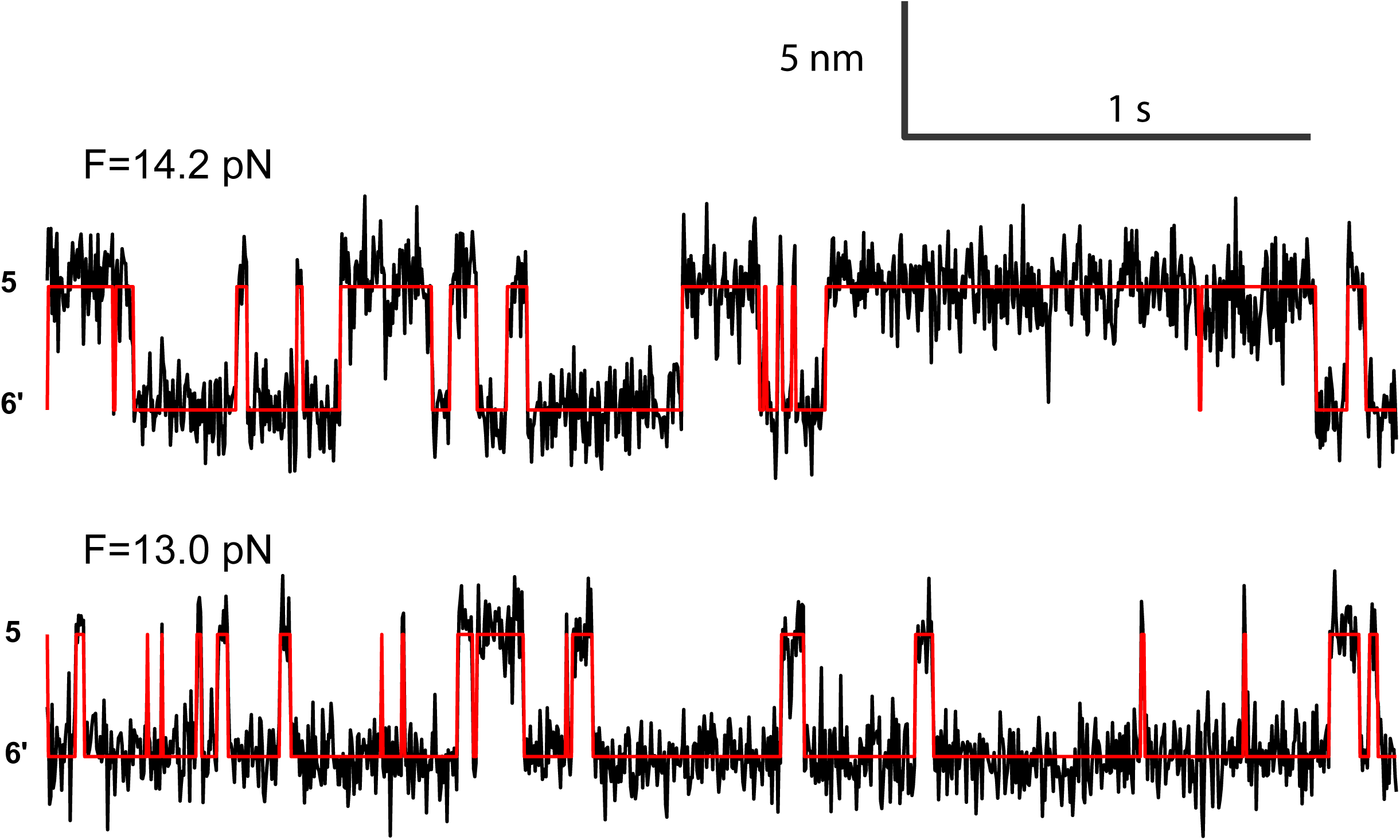
Extension-time trajectories at two constant mean forces (F) showing the opening-closing transition of the syntaxin molecule pulled from the crosslinking site I187C (Figure 2-figure supplement 1). The red curves are idealized state transitions derived from hidden-Markov modeling. State 6’represents the fully closed syntaxin (Figure 2B).

**Figure 2-figure supplement 7.**
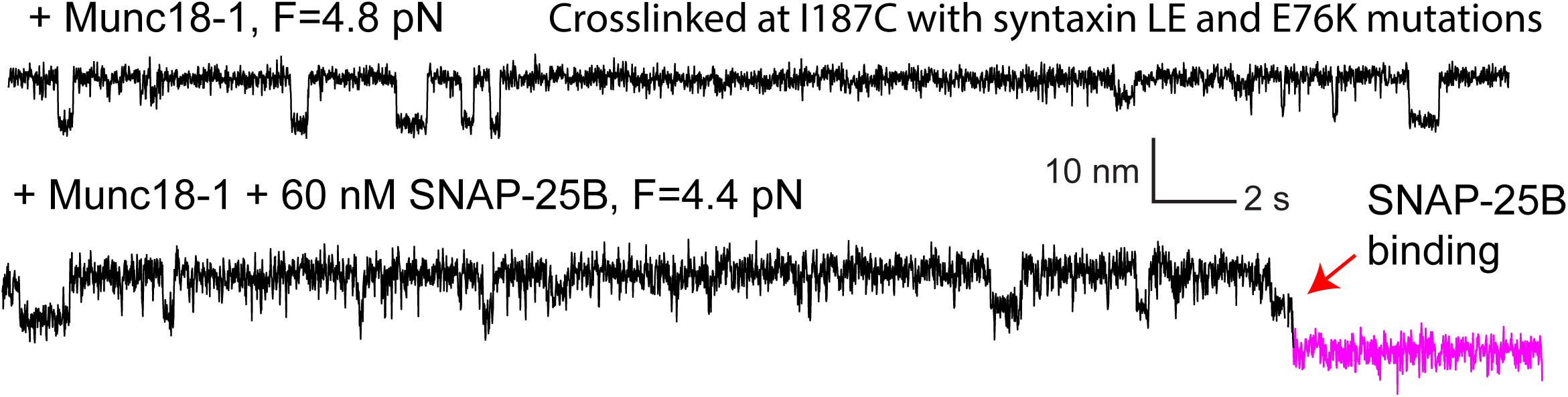
Extension-time trajectories showing conformational transitions of the template complex transition pulled from Site I187C in the absence (top trace) and presence (bottom) of 60 nM SNAP-25B. The unfolding energy of the template complex crosslinked at I187C is estimated to be 4.8 ± 0.3 k_B_T, close to the unfolding energy of 5.2 ±0.1 kBT of the template complex crosslinked at R198C. Furthermore, in the presence of SNAP-25B, the template complex facilitates SNAP-25B binding and SNARE assembly (red arrow).

**Figure 3-figure supplement 1.**
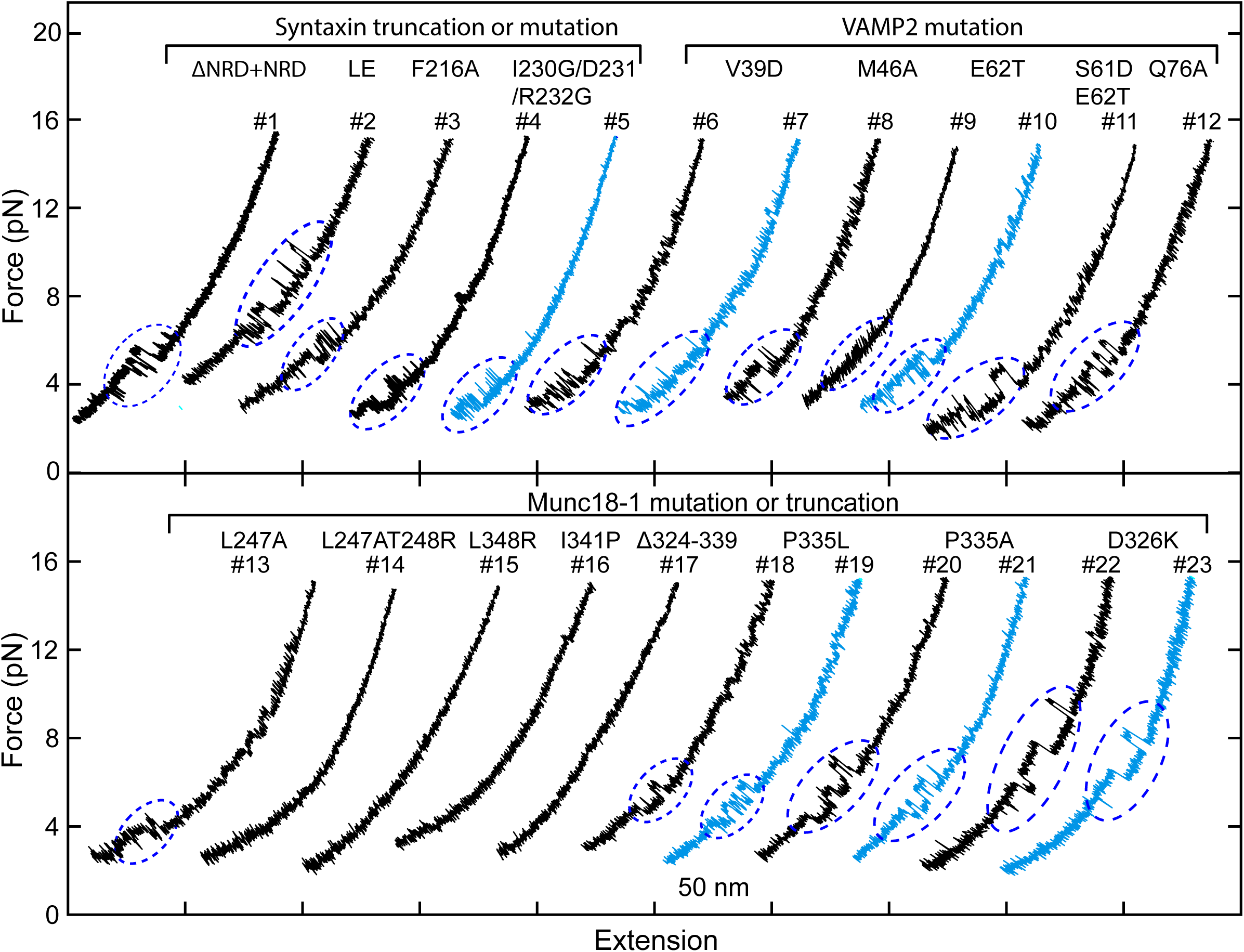
FECs obtained in the presence of 2 µM Munc18-1. Dashed blue ovals mark template complex transitions. Note that the partially closed syntaxin state was abrogated by modifications that are known to destabilize the closed syntaxin, including syntaxin ΔNRD (Burkhardt et al., 2008), the LE mutation (Dulubova et al., 1999; Ma et al., 2011), and Munc18-1 Δ324-339, P335L, and D326K (Munch et al., 2016; Parisotto et al., 2014; Sitarska et al., 2017) (Table 1). For syntaxin LE mutation and I230G/D231/R232G and Munc18-1 P335A and D326K, template complexes generally formed directly from the open syntaxin (Table 1).

**Figure 3-figure supplement 2.**
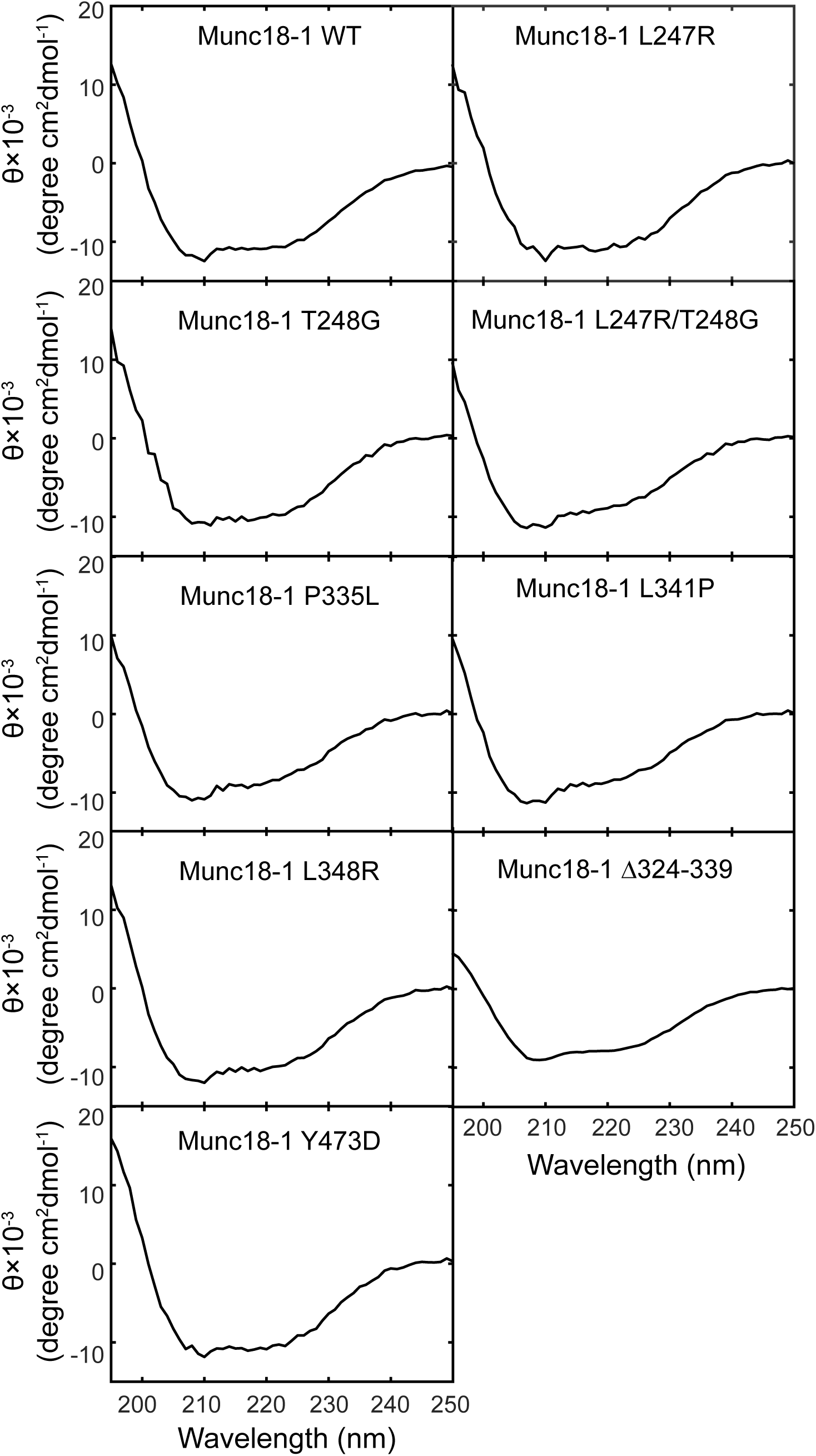
Circular Dichroism (CD) spectra show that mutations barely alter Munc18-1 folding. The CD spectra of Munc18-1 mutants that abolished or weakened the template complex are shown, including Munc18-1 F-pocket mutations L247R, T248G, L247A/T248G, disease-related mutations L341P and P335L, phosphomimetic mutation Y473D, and L348R (Parisotto et al., 2014). All of the mutant proteins displayed CD spectra closely resembling that of wild-type Munc18-1.

**Figure 3-figure supplement 3.**
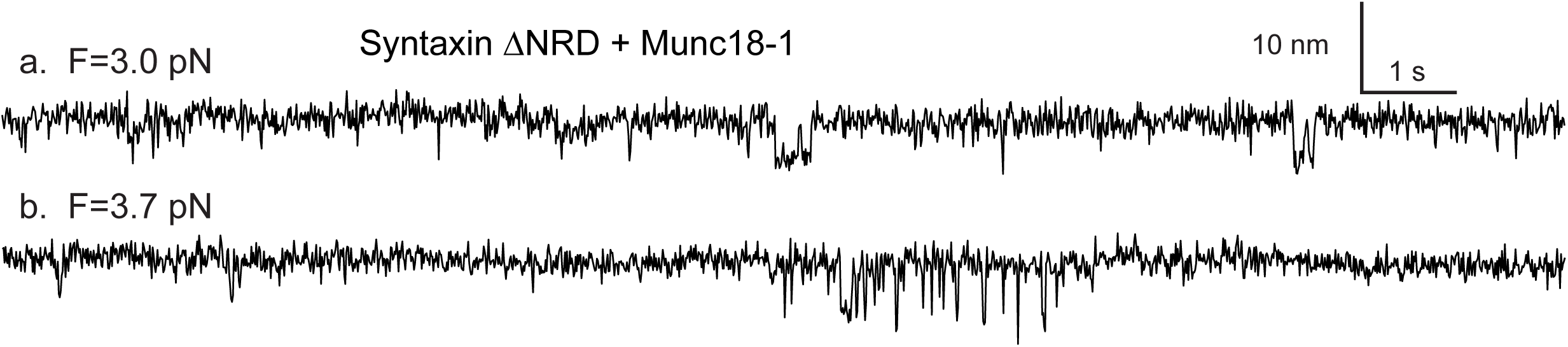
Snapshots of the extension-time trajectories at constant mean forces showing sporadic folding of the template complex in the absence of syntaxin NRD.

**Figure 5-figure supplement 1.**
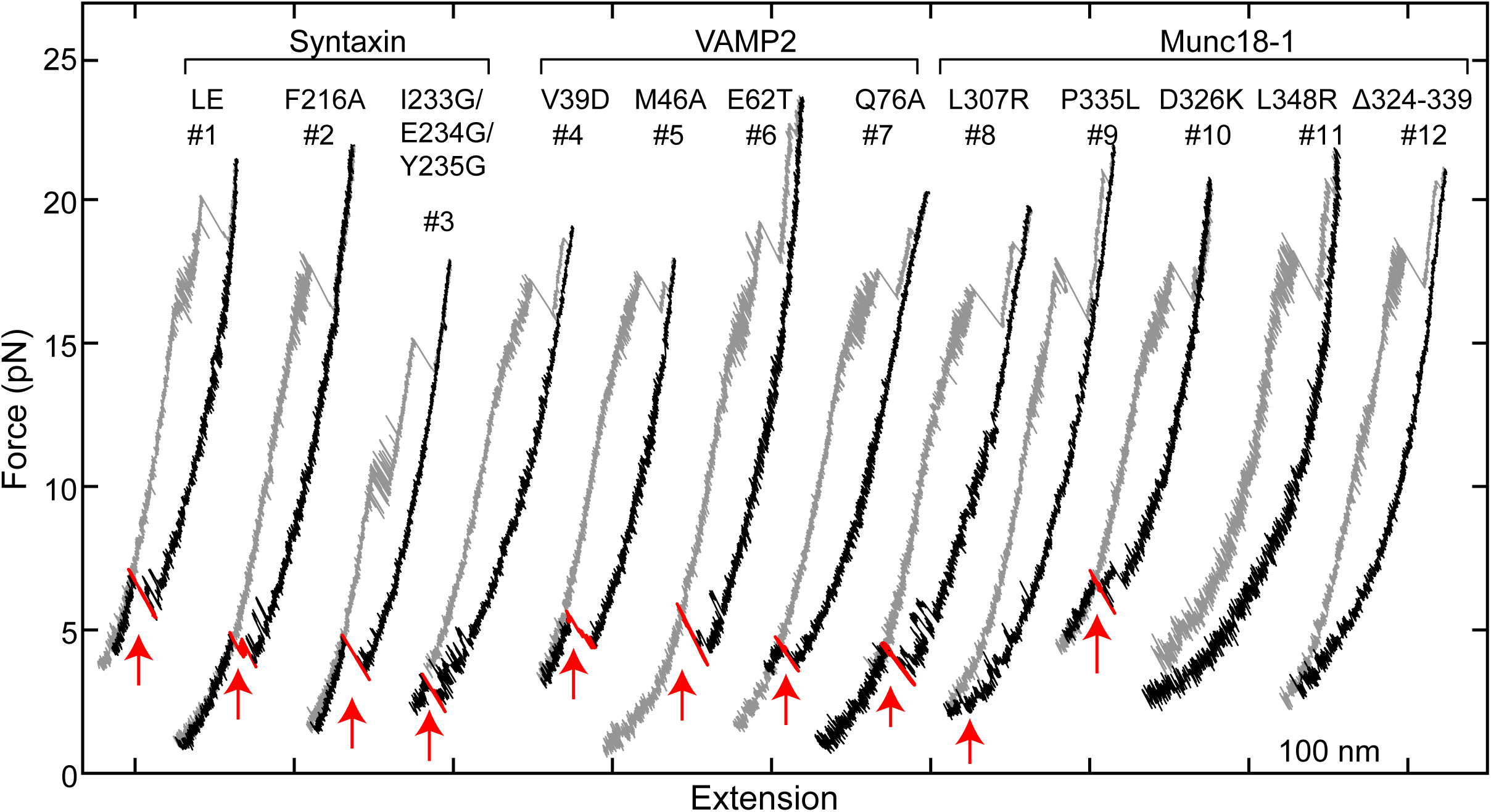
FECs obtained in the presence of 2 µM Munc18-1 and 60 nM SNAP-25B in the solution. Red arrows mark events of SNAP-25B binding and SNARE assembly. Note that syntaxin +2 layer mutation I233G/E234G/Y235G significantly weakens the CTD zippering.

**Figure 5-figure supplement 2.**
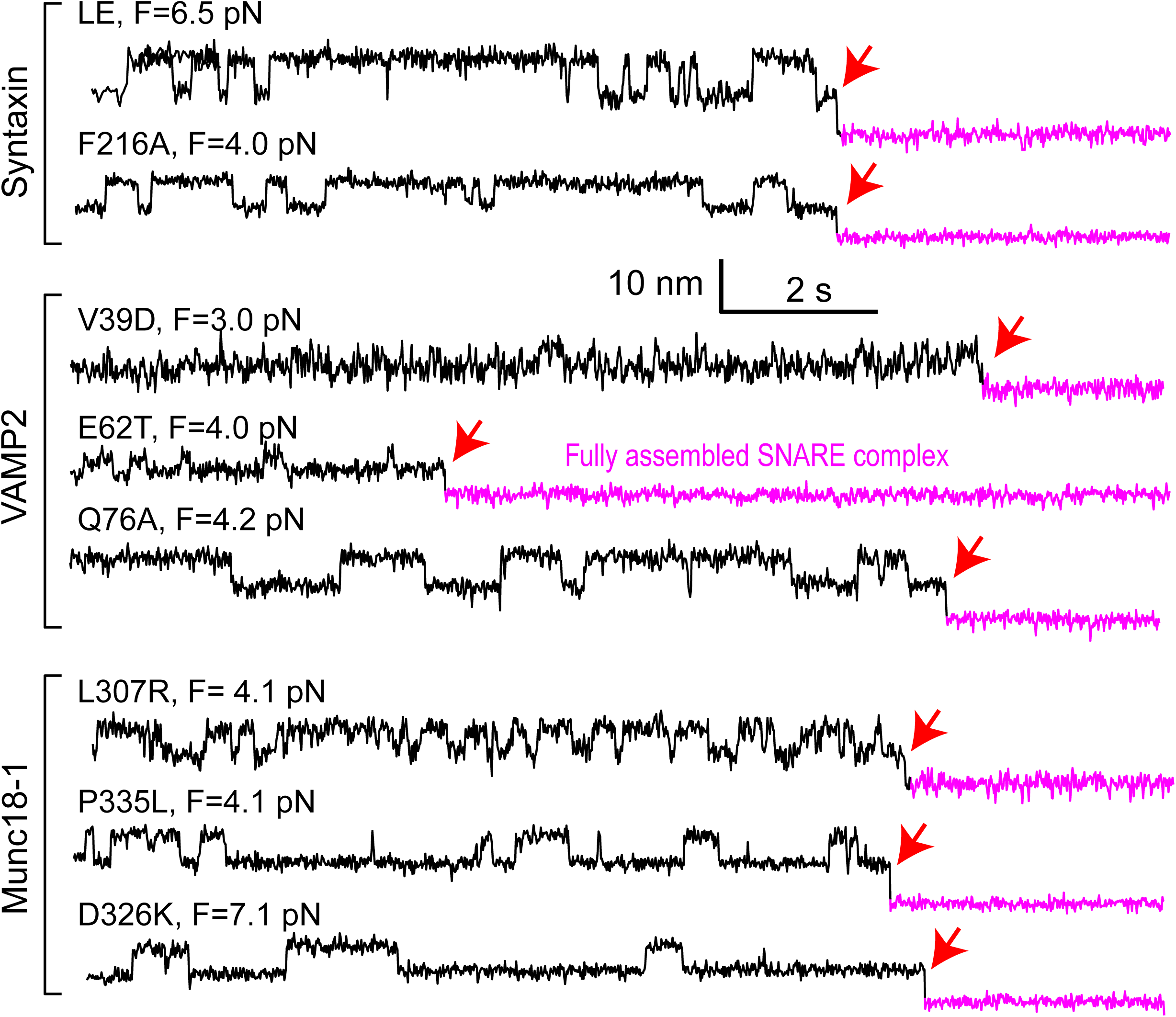
Extension-time trajectories at constant mean forces (F) exhibiting reversible folding and unfolding transitions of the mutant template complexes and irreversible SNAP-25 binding (indicated by red arrows).

**Figure 8-figure supplement 1.**
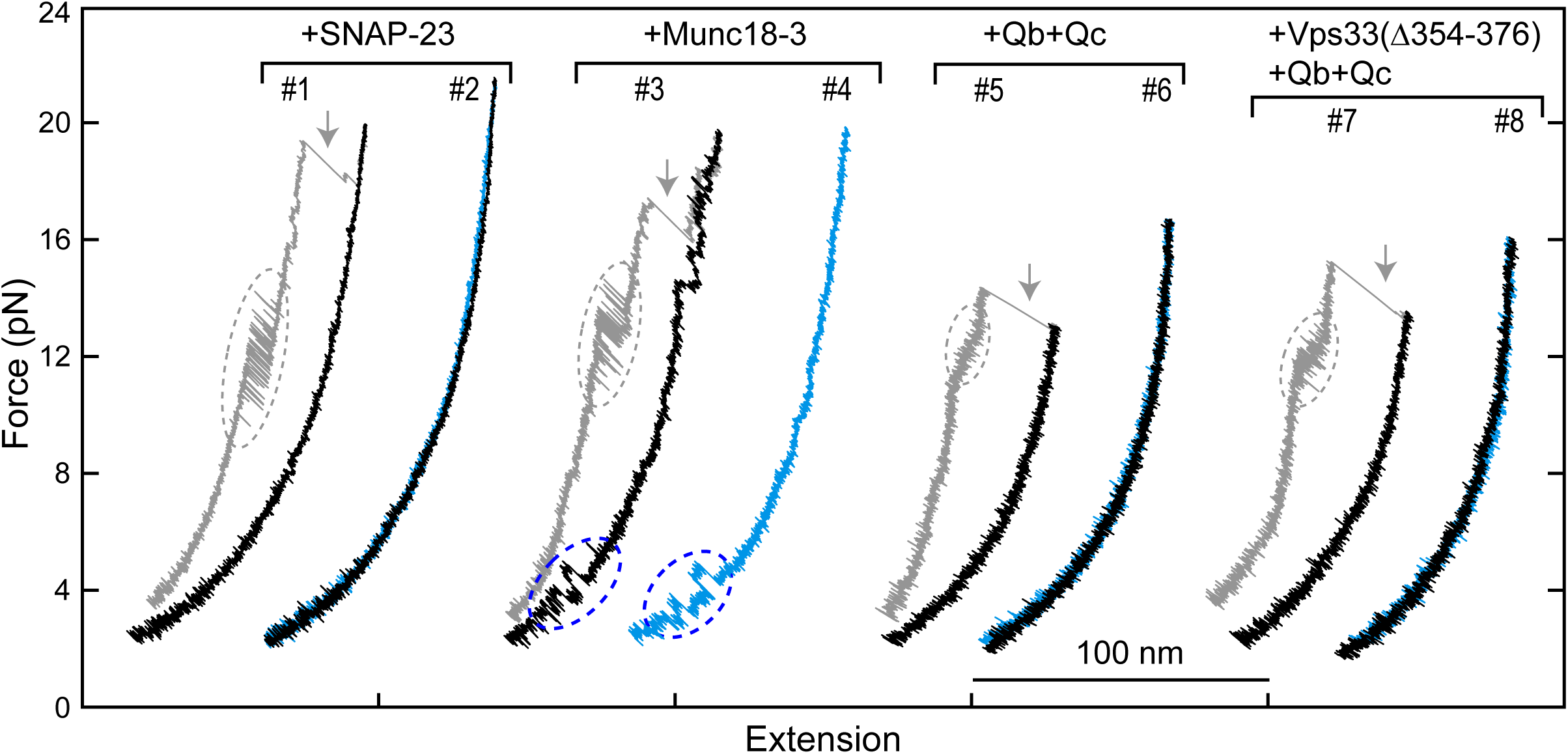
FECs obtained by pulling and relaxing a single syntaxin-4-VAMP2 conjugate (#1-4) or Vam3-Nyv1 conjugate (#5-8) in the presence of the indicated protein or proteins. SNARE CTD transitions and template complex transitions are marked by gray and blue ovals, respectively. Gray arrows indicate SNARE unzipping. Vps33(Δ354-376) is analogous to Munc18-1 Δ324-339 and is inactive in vivo and in vitro (Baker et al., 2015).

**Figure 8-figure supplement 2.**
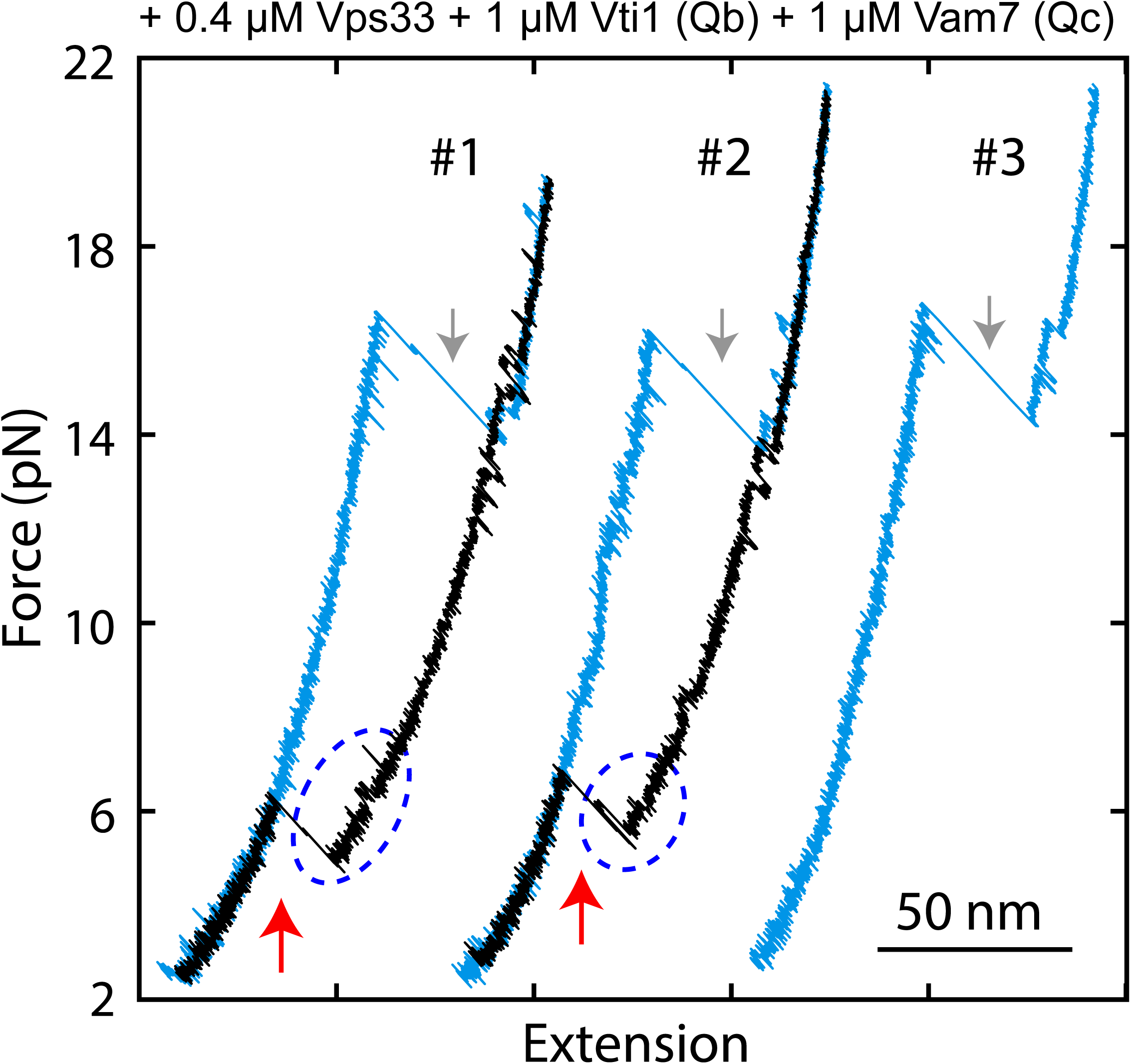
FECs displaying Vps33-catalyzed vacuolar SNARE assembly, marked by red arrows. Red arrows indicate SNARE unzipping. Template complex transitions are marked by blue ovals.

**Figure 8-figure supplement 3.**
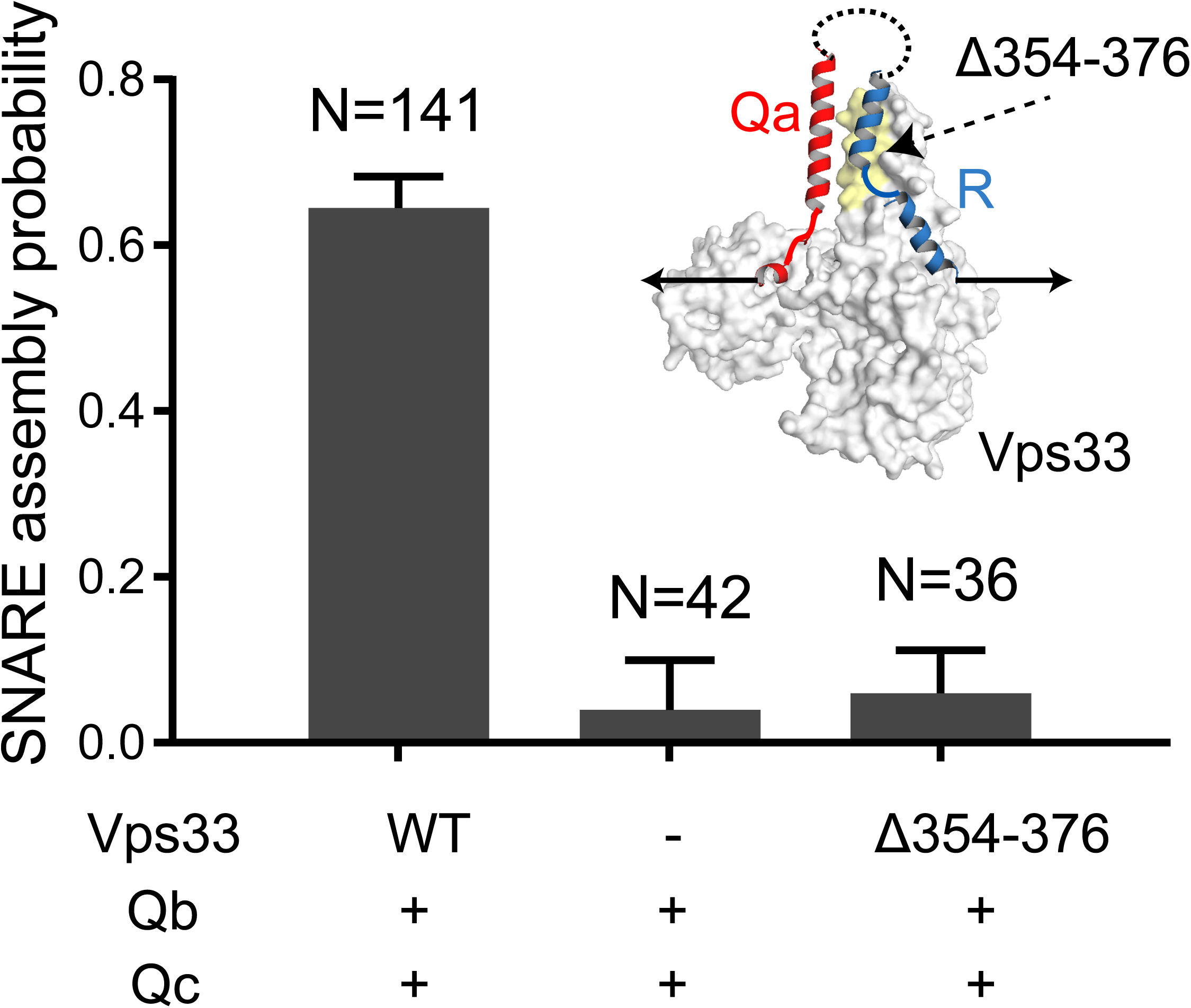
Probabilities of SNARE assembly per relaxation under different conditions. The insert shows the pulling direction and the region of the Vps33 truncation (yellow).

## References

Baker, R.W., and Hughson, F.M. (2016). Chaperoning SNARE assembly and disassembly. Nat Rev Mol Cell Biol 17, 465–479.

Baker, R.W., Jeffrey, P.D., Zick, M., Phillips, B.P., Wickner, W.T., and Hughson, F.M. (2015). A direct role for the Sec1/Munc18-family protein Vps33 as a template for SNARE assembly. Science 349, 1111–1114.

Bao, H., Das, D., Courtney, N.A., Jiang, Y., Briguglio, J.S., Lou, X., Roston, D., Cui, Q., Chanda, B., and Chapman, E.R. (2018). Dynamics and number of trans-SNARE complexes determine nascent fusion pore properties. Nature 554, 260–263.

Brunger, A.T. (2005). Structure and function of SNARE and SNARE-interacting proteins. Q. Rev. Biophys. 38, 1–47.

Brunger, A.T., Choi, U.B., Lai, Y., Leitz, J., and Zhou, Q.J. (2018). Molecular mechanisms of fast neurotransmitter release. Ann Rev Biophys 47, 469–497.

Bryant, N.J., and Gould, G.W. (2011). SNARE proteins underpin insulin-regulated GLUT4 traffic. Traffic 12, 657–664.

Burkhardt, P., Hattendorf, D.A., Weis, W.I., and Fasshauer, D. (2008). Munc18a controls SNARE assembly through its interaction with the syntaxin N-peptide. EMBO J. 27, 923–933.

Colbert, K.N., Hattendorf, D.A., Weiss, T.M., Burkhardt, P., Fasshauer, D., and Weis, W.I. (2013). Syntaxin1a variants lacking an N-peptide or bearing the LE mutation bind to Munc18a in a closed conformation. Proc. Natl. Acad. Sci. U.S.A. 110, 12637–12642.

Cote, M., Menager, M.M., Burgess, A., Mahlaoui, N., Picard, C., Schaffner, C., Al-Manjomi, F., Al-Harbi, M., Alangari, A., Le Deist, F., et al. (2009). Munc18-2 deficiency causes familial hemophagocytic lymphohistiocytosis type 5 and impairs cytotoxic granule exocytosis in patient NK cells. J. Clin. Invest. 119, 3765–3773.

Dawidowski, D., and Cafiso, D.S. (2016). Munc18-1 and the Syntaxin-1 N terminus regulate open-closed states in a t-SNARE complex. Structure 24, 392–400.

Dulubova, I., Khvotchev, M., Liu, S.Q., Huryeva, I., Sudhof, T.C., and Rizo, J. (2007). Munc18-1 binds directly to the neuronal SNARE complex. Proc. Natl. Acad. Sci. U.S.A. 104, 2697–2702.

Dulubova, I., Sugita, S., Hill, S., Hosaka, M., Fernandez, I., Sudhof, T.C., and Rizo, J. (1999). A conformational switch in syntaxin during exocytosis: role of munc18. EMBO J. 18, 4372–4382.

Dulubova, I., Yamaguchi, T., Wang, Y., Sudhof, T.C., and Rizo, J. (2001). Vam3p structure reveals conserved and divergent properties of syntaxins. Nat. Struct. Biol. 8, 258–264.

Fasshauer, D., Sutton, R.B., Brunger, A.T., and Jahn, R. (1998). Conserved structural features of the synaptic fusion complex: SNARE proteins reclassified as Q- and R-SNAREs. Proc. Natl. Acad. Sci. U.S.A. 95, 15781–15786.

Gao, Y., Zorman, S., Gundersen, G., Xi, Z.Q., Ma, L., Sirinakis, G., Rothman, J.E., and Zhang, Y.L. (2012). Single reconstituted neuronal SNARE complexes zipper in three distinct stages. Science 337, 1340–1343.

Genc, O., Kochubey, O., Toonen, R.F., Verhage, M., and Schneggenburger, R. (2014). Munc18-1 is a dynamically regulated PKC target during short-term enhancement of transmitter release. eLife 3, e01715.

Gerber, S.H., Rah, J.C., Min, S.W., Liu, X.R., de Wit, H., Dulubova, I., Meyer, A.C., Rizo, J., Arancillo, M., Hammer, R.E., et al. (2008). Conformational switch of syntaxin-1 controls synaptic vesicle fusion. Science 321, 1507–1510.

Hu, S.H., Christie, M.P., Saez, N.J., Latham, C.F., Jarrott, R., Lua, L.H.L., Collins, B.M., and Martin, J.L. (2011). Possible roles for Munc18-1 domain 3a and Syntaxin1 N-peptide and C-terminal anchor in SNARE complex formation. Proc. Natl. Acad. Sci. U.S.A. 108, 1040–1045.

Jakhanwal, S., Lee, C.T., Urlaub, H., and Jahn, R. (2017). An activated Q-SNARE/SM protein complex as a possible intermediate in SNARE assembly. EMBO J. 36, 1788–1802.

Jiao, J.Y., Rebane, A.A., Ma, L., and Zhang, Y.L. (2017). Single-molecule protein folding experiments using high-resolution optical tweezers. Methods Mol Biol 1486, 357–390.

Lai, Y., Choi, U.B., Leitz, J., Rhee, H.J., Lee, C., Altas, B., Zhao, M.L., Pfuetzner, R.A., Wang, A.L., Brose, N., et al. (2017). Molecular mechanisms of synaptic vesicle priming by Munc13 and Munc18. Neuron 95, 591–607.

Ma, C., Li, W., Xu, Y., and Rizo, J. (2011). Munc13 mediates the transition from the closed syntaxin-Munc18 complex to the SNARE complex. Nat Struct Mol Biol 18, 542–549.

Ma, C., Su, L.J., Seven, A.B., Xu, Y.B., and Rizo, J. (2013). Reconstitution of the vital functions of Munc18 and Munc13 in neurotransmitter release. Science 339, 421–425.

Ma, L., Rebane, A.A., Yang, G., Xi, Z., Kang, Y., Gao, Y., and Zhang, Y.L. (2015). Munc18-1-regulated stage-wise SNARE assembly underlying synaptic exocytosis. eLife 4, e09580.

McKinney, S.A., Joo, C., and Ha, T. (2006). Analysis of single-molecule FRET trajectories using hidden Markov modeling. Biophys. J. 91, 1941–1951.

Meijer, M., Burkhardt, P., de Wit, H., Toonen, R.F., Fasshauer, D., and Verhage, M. (2012). Munc18-1 mutations that strongly impair SNARE-complex binding support normal synaptic transmission. EMBO J. 31, 2156–2168.

Meijer, M., Dorr, B., Lammertse, H.C.A., Blithikioti, C., van Weering, J.R.T., Toonen, R.F.G., Sollner, T.H., and Verhage, M. (2018). Tyrosine phosphorylation of Munc18-1 inhibits synaptic transmission by preventing SNARE assembly. EMBO J. 37, 300–320.

Misura, K.M.S., Scheller, R.H., and Weis, W.I. (2000). Three-dimensional structure of the neuronal-Sec1-syntaxin 1a complex. Nature 404, 355–362.

Mohrmann, R., de Wit, H., Verhage, M., Neher, E., and Sorensen, J.B. (2010). Fast vesicle fusion in living cells requires at least three SNARE complexes. Science 330, 502–505.

Morey, C., Kienle, C.N., Klopper, T.H., Burkhardt, P., and Fasshauer, D. (2017). Evidence for a conserved inhibitory binding mode between the membrane fusion assembly factors Munc18 and syntaxin in animals. J. Biol. Chem. 292, 20449–20460.

Munch, A.S., Kedar, G.H., van Weering, J.R.T., Vazquez-Sanchez, S., He, E.Q., Andre, T., Braun, T., Sollner, T.H., Verhage, M., and Sorensen, J.B. (2016). Extension of Helix 12 in Munc18-1 induces vesicle priming. J. Neurosci. 36, 6881–6891.

Parisotto, D., Pfau, M., Scheutzow, A., Wild, K., Mayer, M.P., Malsam, J., Sinning, I., and Sollner, T.H. (2014). An extended helical conformation in domain 3a of Munc18-1 provides a template for SNARE (soluble N-ethylmaleimidesensitive factor attachment protein receptor) complex assembly. J. Biol. Chem. 289, 9639–9650.

Pobbati, A.V., Stein, A., and Fasshauer, D. (2006). N-to C-terminal SNARE complex assembly promotes rapid membrane fusion. Science 313, 673–676.

Rebane, A.A., Ma, L., and Zhang, Y.L. (2016). Structure-based derivation of protein folding intermediates and energies from optical tweezers. Biophys J 110, 441–454.

Rebane, A.A., Wang, B., Ma, L., Qu, H., Coleman, J., Krishnakumar, S.S., Rothman, J.E., and Zhang, Y.L. (2018). Two disease-causing SNAP-25B mutations selectively impair SNARE C-terminal assembly. J. Mol. Biol. 430, 479–490.

Richmond, J.E., Weimer, R.M., and Jorgensen, E.M. (2001). An open form of syntaxin bypasses the requirement for UNC-13 in vesicle priming. Nature 412, 338–341.

Rizo, J., and Sudhof, T.C. (2012). The membrane fusion enigma: SNAREs, Sec1/Munc18 proteins, and their accomplices-guilty as charged? Annu. Rev. Cell. Dev. Biol. 28, 279–308.

Shen, J.S., Rathore, S.S., Khandan, L., and Rothman, J.E. (2010). SNARE bundle and syntaxin N-peptide constitute a minimal complement for Munc18-1 activation of membrane fusion. J. Cell Biol. 190, 55–63.

Shen, J.S., Tareste, D.C., Paumet, F., Rothman, J.E., and Melia, T.J. (2007). Selective activation of cognate SNAREpins by Sec1/Munc18 proteins. Cell 128, 183–195.

Sitarska, E., Xu, J.J., Park, S., Liu, X.X., Quade, B., Stepien, K., Sugita, K., Brautigam, C.A., Sugita, S., and Rizo, J. (2017). Autoinhibition of Munc18-1 modulates synaptobrevin binding and helps to enable Munc13-dependent regulation of membrane fusion. eLife 6, e24278.

Sollner, T., Whiteheart, S.W., Brunner, M., Erdjument-Bromage, H., Geromanos, S., Tempst, P., and Rothman, J.E. (1993). SNAP receptors implicated in vesicle targeting and fusion. Nature 362, 318–324.

Stamberger, H., Nikanorova, M., Willemsen, M.H., Accorsi, P., Angriman, M., Baier, H., Benkel-Herrenbrueck, I., Benoit, V., Budetta, M., Caliebe, A., et al. (2016). STXBP1 encephalopathy: A neurodevelopmental disorder including epilepsy. Neurology 86, 954–962.

Sudhof, T.C., and Rothman, J.E. (2009). Membrane fusion: grappling with SNARE and SM proteins. Science 323, 474–477.

Sutton, R.B., Fasshauer, D., Jahn, R., and Brunger, A.T. (1998). Crystal structure of a SNARE complex involved in synaptic exocytosis at 2.4 angstrom resolution. Nature 395, 347–353.

Verhage, M., Maia, A.S., Plomp, J.J., Brussaard, A.B., Heeroma, J.H., Vermeer, H., Toonen, R.F., Hammer, R.E., van den Berg, T.K., Missler, M., et al. (2000). Synaptic assembly of the brain in the absence of neurotransmitter secretion. Science 287, 864–869.

Walter, A.M., Wiederhold, K., Bruns, D., Fasshauer, D., and Sorensen, J.B. (2010). Synaptobrevin N-terminally bound to syntaxin-SNAP-25 defines the primed vesicle state in regulated exocytosis. J. Cell Biol. 188, 401–413.

Wang, S., Choi, U.B., Gong, J.H., Yang, X.Y., Li, Y., Wang, A.L., Yang, X.F., Brunger, A.T., and Ma, C. (2017). Conformational change of syntaxin linker region induced by Munc13s initiates SNARE complex formation in synaptic exocytosis. EMBO J. 36, 816–829.

Weber, T., Zemelman, B.V., McNew, J.A., Westermann, B., Gmachl, M., Parlati, F., Sollner, T.H., and Rothman, J.E. (1998). SNAREpins: Minimal machinery for membrane fusion. Cell 92, 759–772.

Wickner, W. (2010). Membrane fusion: Five lipids, four SNAREs, three chaperones, two nucleotides, and a Rab, all dancing in a ring on yeast vacuoles. Annu Rev Cell Dev Bi 26, 115–136.

Yang, X.Y., Wang, S., Sheng, Y., Zhang, M.S., Zou, W.J., Wu, L.J., Kang, L.J., Rizo, J., Zhang, R.G., Xu, T., et al. (2015). Syntaxin opening by the MUN domain underlies the function of Munc13 in synaptic-vesicle priming. Nat. Struct. Mol. Biol. 22, 547–754.

Zhang, X.M., Rebane, A.A., Ma, L., Li, F., Jiao, J., Qu, H., Pincet, F., Rothman, J.E., and Zhang, Y.L. (2016a). Stability, folding dynamics, and long-range conformational transition of the synaptic t-SNARE complex. Proc. Natl. Acad. Sci. U.S.A. 113, E8031–E8040.

Zhang, Y., Diao, J., Colbert, K.N., Lai, Y., Pfuetzner, R.A., Padolina, M.S., Vivona, S., Ressl, S., Cipriano, D.J., Choi, U.B., et al. (2015). Munc18a does not alter fusion rates mediated by neuronal snares, synaptotagmin, and complexin. J Biol Chem 290, 10518–10534.

Zhang, Y.L., Jiao, J., and Rebane, A.A. (2016b). Hidden Markov modeling with detailed balance and its application to single protein folding Biophys J 111, 2110–2124.

Zhou, P., Pang, Z.P.P., Yang, X.F., Zhang, Y.S., Rosenmund, C., Bacaj, T., and Sudhof, T.C. (2013). Syntaxin-1 N-peptide and Habc-domain perform distinct essential functions in synaptic vesicle fusion. EMBO J. 32, 159–171.

Zorman, S., Rebane, A.A., Ma, L., Yang, G.C., Molski, M.A., Coleman, J., Pincet, F., Rothman, J.E., and Zhang, Y.L. (2014). Common intermediates and kinetics, but different energetics, in the assembly of SNARE proteins. eLife 3, e03348.

